# Reducing functionally defective old HSCs alleviates aging-related phenotypes in old recipient mice

**DOI:** 10.1101/2024.11.09.622774

**Authors:** Yuting Wang, Wenhao Zhang, Chao Zhang, Hoang Tran Van, Takashi Seino, Yi Zhang

## Abstract

Aging is a process accompanied by functional decline in tissues and organs with great social and medical consequences. Developing effective anti-aging strategies is of great significance. In this study, we demonstrated that transplantation of young hematopoietic stem cells (HSCs) into old mice can mitigate aging phenotypes, underscoring the crucial role of HSCs in the aging process. Through comprehensive molecular and functional analyses, we identified a subset of HSCs in aged mice that exhibit “younger” molecular profiles and functions, marked by low levels of CD150 expression. Mechanistically, CD150^low^ HSCs from old mice can effectively differentiate into downstream lineage cells but not their CD150^high^ counterparts. Notably, transplantation of old CD150^low^ HSCs attenuates aging phenotypes and prolongs lifespan of elderly mice compared to those transplanted with unselected or CD150^high^ HSCs. Importantly, reducing the dysfunctional CD150^high^ HSCs can alleviate aging phenotypes in old recipient mice. Thus, our study demonstrates the presence of “younger” HSCs in old mice, and aging-associated functional decline can be mitigated by reducing dysfunctional HSCs.

## Introduction

The aging process is marked by a functional decline across various tissues and organs,^1–3^ which significantly increases the risk for a multitude of chronic diseases.^4^ Previous studies have shown that transfusion of young blood or its components into aged mice can have a rejuvenation effect,^5–10^ supporting an important role of the hematopoietic system in whole body aging. Hematopoietic stem cells (HSCs), the source of all hematopoietic cell types, undergo profound functional changes with age, including increased prevalence of clonal hematopoiesis, a shift toward myeloid-biased differentiation, and a diminished capacity for blood regeneration.^11–15^ Recent studies have shown that replacing HSCs of old mice with those of young mice can improve health and extend lifespan of old recipient mice.^16–18^ Conversely, transplanting aged HSCs into young mice can accelerate immune system aging and reduce their lifespan.^19, 20^ These studies suggest that HSCs could serve as promising targets for aging-related intervention.

Previous studies have established that intrinsic alterations of HSCs with aging are associated with their functional decline, including increased DNA damage, mitochondrial degeneration, diminished asymmetric distribution of CDC42, altered H4K16 acetylation during mitosis, and other epigenetic modifications.^21–27^ Interestingly, the HSC population exhibits intrinsic heterogeneity in both young and old mice, distinguishable by lymphoid-biased, balanced, and myeloid-biased subtypes based on surface markers and lineage predispositions.^28–36^ Notably, a fraction of HSCs can preserve long-term dormancy and functionality during aging.^37–39^ However, the changes of HSC heterogeneity during aging and their contribution to systemic aging remains to be comprehensively characterized.

In this study, we demonstrate that transplantation of young HSCs can alleviate aging phenotypes in old mice. scRNA-seq reveals an increased aging heterogeneity in old HSCs. Systematic molecular and functional characterization indicate the presence of “younger” and functionally defective “older” subsets of HSCs in old mice, marked by differential expression levels of CD150. Mechanistically, old CD150^high^ HSCs exhibit differentiation defects compared to those of the CD150^low^ HSCs. Notably, transplantation of the “younger” subset of HSCs into old recipient mice can attenuate aging phenotypes, including decreased epigenetic age and extended lifespan of old recipient mice compared with those that received old un-selected HSCs or CD150^high^ HSCs. Importantly, reducing the dysfunctional CD150^high^ HSCs can attenuate aging phenotypes in old recipient mice, highlighting that the removal of defective CD150^high^ HSCs from old mice could be a potential strategy for rejuvenation.

## Results

### Transplantation of young HSCs alleviates aging phenotypes in old recipient mice

To directly evaluate the functional differences between young and old HSCs in vivo, we sorted long-term (LT)-HSCs [lineage^-^c-Kit^+^Sca-1^+^ (LSK) CD48^-^CD34^-^CD150^+^] (referred to as HSCs) from young and old mice (**Supplementary information, Fig. S1a**). Consistent with previous reports,^12, 13, 40^ a significant increase in the number of HSCs in old mice was observed (**Supplementary information, Fig. S1b**). Serial transplantation of both young and old HSCs into lethally irradiated young recipient mice (**Fig. 1a**) revealed a notable decline in the repopulation capacity and biased differentiation of old HSCs compared to their young counterparts (**Fig. 1b; Supplementary information, Fig. S1c-f**). These results are consistent with previous studies,^12, 20^ and confirm the age-related intrinsic functional decline of HSCs.

**Fig. 1.**
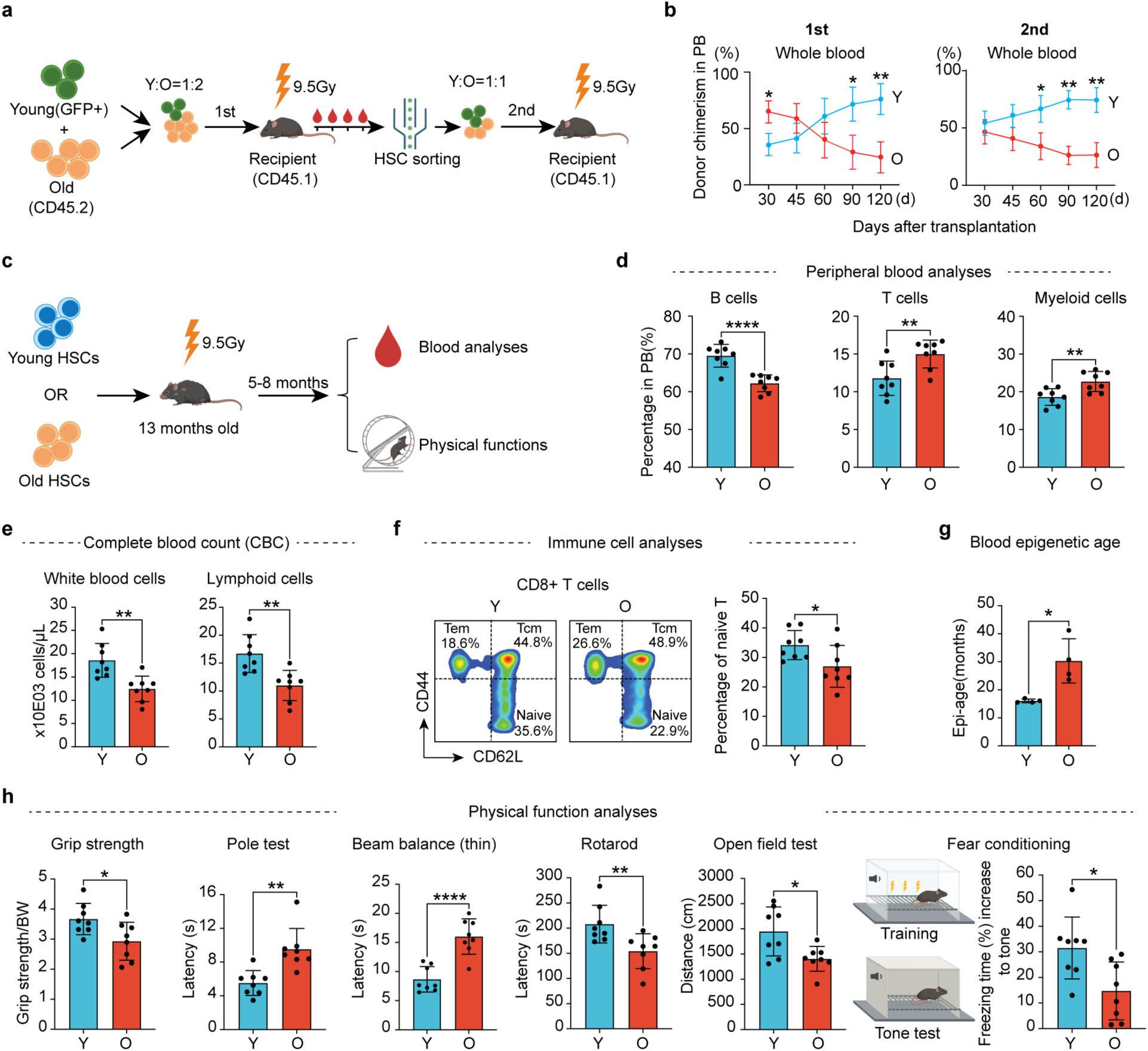
Transplantation of young HSCs alleviates aging phenotypes in old recipient mice. **a** Diagram illustration of the competitive young (2-3 months) and old (22-24 months) HSC transplantation experiment. For the first transplantation, the ratio of young to old HSCs was 1:2 (500 young and 1,000 old) while the ratio was 1:1 (1,000 young and 1,000 old) for the second transplantation. **b** Peripheral blood chimerism of donor HSCs at different times after the 1st and 2nd transplantation. *n* = 6 for the first and *n* = 3 for the second transplantation. **c** Diagram showing the experimental design of individual transplantation of 1,000 young (3 months) or 1,000 old (22-24 months) HSCs into middle-aged recipients (13-month-old). Five months after the transplantation, a series of hematopoietic and physical tests were performed. **d** Bar graph showing the percentage of B, T and myeloid cells in the peripheral blood (PB) of recipient mice, *n* = 8. **e** Bar graph showing the absolute number of blood cells in the PB of recipient mice, *n* = 8. **f** FACS plot and bar plot showing the percentage of naïve T cells in CD8+ T cells from recipient mice, Naïve T cells (CD44^low^, CD62L^high^), Tcm (CD44^high^, CD62L^high^) and Tem (CD44^high^, CD62L^low^), *n* = 8. **g** Bar plot showing the epigenetic age of blood from recipient mice, *n* = 4. **h** Physical tests of recipient mice that received young or old HSCs. Muscle strength, motor coordination, endurance and brain function were assessed, *n* = 8. Mean ± SD, student t test, \**P*<0.05, ** *P*<0.01, **** *P*<0.0001. The graphic of the mouse and equipment in **a**, **c** and **h** were created with BioRender.

To analyze the contribution of HSCs to systemic aging, we transplanted young or old HSCs into irradiated middle-aged mice (13 months). Five months after transplantation, we evaluated the aging status of the recipient mice by performing blood analyses and physical tests (**Fig. 1c**). Previous studies have shown that the blood cell composition changes with aging, characterized by an increase in myeloid cells and a corresponding decrease in lymphoid cells.^41, 42^ Notably, mice transplanted with young HSCs exhibited a more youthful blood cell composition, characterized by increased B cell and reduced T cell and myeloid cell proportions, as well as an increase in overall white blood cells and lymphoid counts, compared to the recipients that received old HSCs (**Fig. 1d, e**). This suggests a crucial role of HSCs in systematic hematopoietic aging.

The ratio of naïve T cells serves as a critical indicator of immune function, which declines with aging.^43^ Conversely, central memory T cells (Tcm) and effector memory T cells (Tem) exhibit an increase with aging,^44^ which were confirmed by our study (**Supplementary information, Fig. S2**). Notably, the old mice that received young HSCs have more naïve T cells in both CD4 and CD8 T cells (**Fig. 1f; Supplementary information, Fig. S3a, b**), indicating an improved immune system after receiving young HSCs transplantation.

The DNA methylation clock is widely used as an indicator of aging.^45^ Importantly, old mice that received young HSCs have a younger epigenetic age in blood compared to those that received old HSCs (**Fig. 1g**). Consistently, old mice that received young HSCs exhibited improved physical functions when compared to those that received aged HSCs, including muscle strength, motor coordination, locomotor activity and cognitive functions (**Fig. 1h**), without significant difference in body weight and spatial memory (**Supplementary information, Fig. S3c-e**). Collectively, these results demonstrate that transplantation of young HSCs to old mice can alleviate aging-related phenotypes.

### Single cell RNA-seq reveals a “younger” HSC subpopulation in old mice

The above results suggest that HSCs from old mice are functionally defective compared to those from young mice. To understand the defects, we performed both bulk RNA-seq and scRNA-seq on young and old HSCs (**Fig. 2a**), which revealed a clear transcriptomic difference between young and old HSCs (**Fig. 2b; Supplementary information, Fig. S4a**). Bulk RNA-seq revealed 332 up-regulated genes in old HSCs, including previously annotated HSC aging marker genes,^46^ including *Clu*, *Selp*, *Mt1* and *Ramp2* (**Supplementary information, Fig. S4a; Supplementary information, Table S1**). These genes were defined as HSC aging marker genes for downstream analysis. Interestingly, unsupervised clustering analysis of scRNA-seq revealed that while the quiescent young HSCs are largely uniform, the quiescent old HSCs can be further divided into 3 clusters (q1-3) (**Fig. 2c, d**). This result indicates that quiescent old HSCs are transcriptionally more heterogeneous compared to young HSCs.

**Fig. 2.**
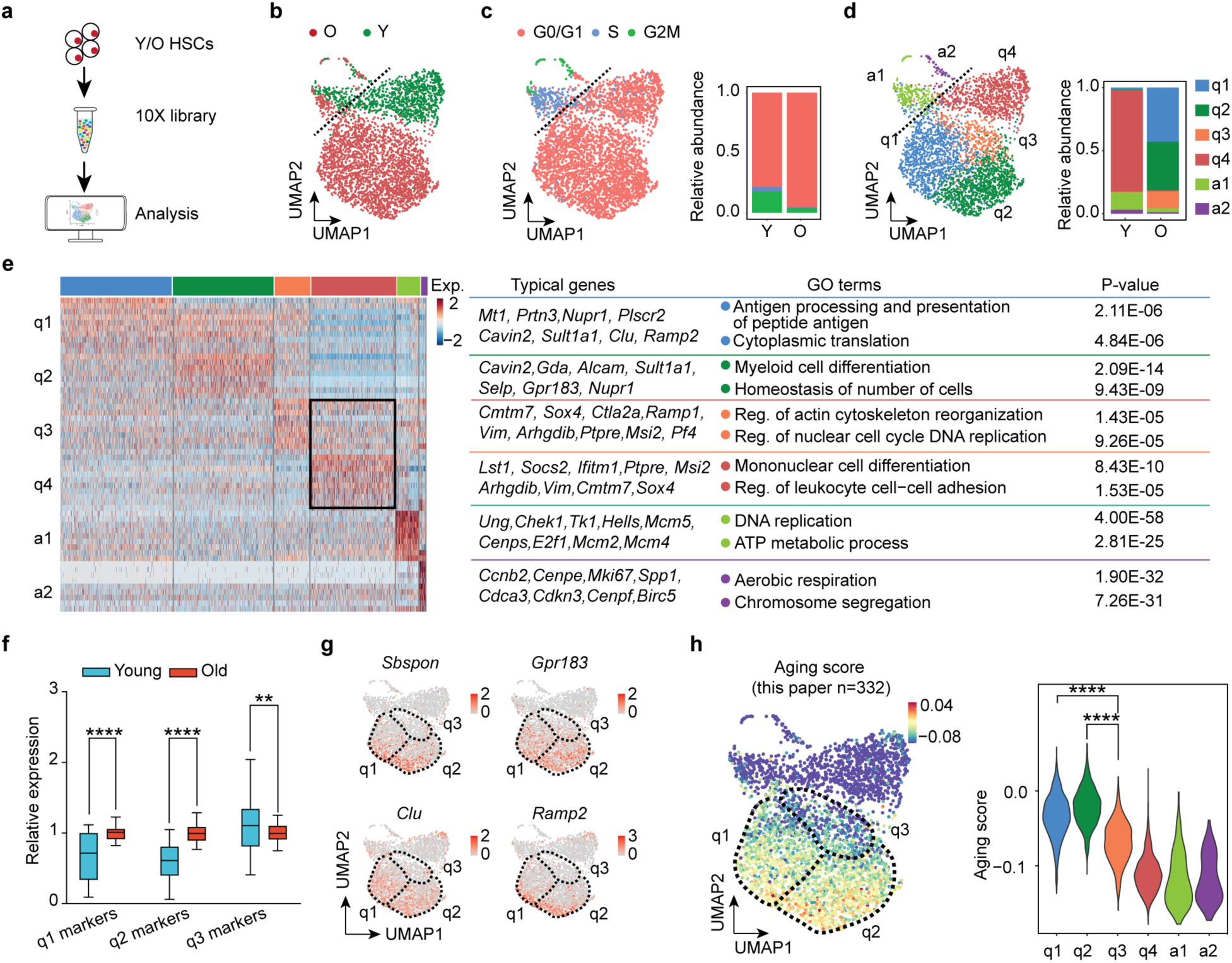
scRNA-seq reveals increased heterogeneity of old HSCs. **a** Workflow of 10X scRNA-seq of young and old HSCs. The HSCs were sorted from young (2-3 months) and old (23 months) mice. **b** UMAP plot showing the distribution of young and old HSCs based on scRNA-seq. Clear separation of young and old HSCs indicates transcriptional changes of HSCs during aging. **c** Cell cycle phase analysis of young and old HSCs based on scRNA-seq. The S-phase and G2/M-phase marker genes are from Seurat package (V4.0.2). The cell cycle phase is determined by the relative expression levels of these marker genes. If neither S-phase nor G2/M phase genes are expressed, they are classified as G0/G1 phase. **d** UMAP plot showing unsupervised clustering of young and old HSCs. In total, 6 clusters were identified with two clusters (a1 and a2) representing active, and four clusters (q1-q4) representing quiescent cells. **e** Heatmap showing the cluster-specific marker genes expression across the 6 clusters and their enriched GO terms. Differentially expressed genes with min.pct = 0.25, logfc.threshold = 0.25 among the 6 clusters were used to generate the heatmap. Well-known HSC aging related genes in clusters q1 and q2 are highlighted. Commonly identified marker genes in clusters q3 and q4 were also highlighted. **f** Box plot showing relative expression of marker genes of q1, q2 and q3 in young and old HSCs via analysis of bulk RNA-seq. The expression level was normalized to the average expression level of the old. Plot shows the mean and 5-95 percentile. Two-sided unpaired Wilcoxon test. **g** UMAP presentation of well-known HSC aging marker genes, *Sbspon*, *Gpr183*, *Clu* and *Ramp2* in each of the single cells. **h** UMAP plot and violin plot showing the calculated aging score of single cells of different clusters based on the HSC aging genes identified in bulk RNA-seq. In total, 332 up-regulated genes in old HSCs were used for aging score calculation. Two-sided unpaired Wilcoxon test. \*\**P*<0.01, \*\*\*\**P*<0.0001.

We then identified the marker genes for different clusters (**Fig. 2e; Supplementary information, Table S2**). Intriguingly, although q1-q3 are all from old HSCs, the known aging marker genes, such as *Mt1*, *Nupr1, Cavin2, Clu, Ramp2, Alcam, Selp* and *Gpr183*, are highly expressed in q1 and q2 clusters, but not in q3 cluster (**Fig. 2e**). In contrast, q3 and q4 (young HSCs) clusters share highly expressed genes, despite the q3 clusters originating from old mice. GO analysis of q3 marker genes revealed an enrichment of cell proliferation related pathways (**Fig. 2e**), which were also enriched in the young HSCs (**Supplementary information, Fig. S4b**, aging-down). We further calculated the relative expression levels of q1, q2, and q3 marker genes in old and young HSCs, and found that the q1 and q2 marker genes are expressed significantly higher in old HSCs, while q3 marker genes are higher in young HSCs (**Fig. 2f**), supporting a younger transcriptome of q3 HSCs.

To gain further support for the notion that a subset of HSCs from old mice are “younger” in their transcriptome, we analyzed the UMAP distribution of some well-known HSC aging marker genes (*Sbspon*, *Gpr183*, *Clu*, *Ramp2*) ^46^ and found that they are expressed at a higher level in q1 and q2 compared to that in q3 (**Fig. 2g**). In contrast, the young HSC marker genes, such as *Rnase6* and *Arhgap30*, exhibited an opposite expression pattern (**Supplementary information, Fig. S4c**). To quantify the aging heterogeneity of old HSCs, we calculated the aging score using the 332 HSC aging marker genes (**Supplementary information, Table S1**) or top 100 reported HSCs aging genes (**Supplementary information, Table S3**).^46^ Both analyses indicate that cluster q3 has a lower aging score than q1 and q2, despite all cells originating from the same old mice (**Fig. 2h; Supplementary information, Fig. S4d**). Collectively, comparative scRNA-seq revealed increased aging heterogeneity in old HSCs and a subset of old HSCs have a younger transcriptome.

### CD150 can serve as an aging heterogeneity marker for old HSCs

To better understand the aging heterogeneity of old HSCs and to assess its contribution to overall body aging, a unique cell surface marker that facilitates the separation of “younger” from “older” cells in aged HSCs is needed. To this end, we first identified the top 150 genes showing strong correlation with aging scores in scRNA-seq (**Supplementary information, Fig. S5a**) and compared them with the 332 genes upregulated in old HSCs (**Supplementary information, Fig. S4a**), resulting in 54 shared genes (**Fig. 3a; Supplementary information, Table S4**). We further required membrane location for sorting purposes, which reduced the candidate gene list to 26 (**Fig. 3a**). Furthermore, the potential marker genes should have lower expression levels in q3 than in q1 and q2, which further narrowed the list to 7 (**Supplementary information, Fig. S5b**). Considering the availability and specificity of the antibodies, CD150 (*Slamf1*) was selected as the marker for aging heterogeneity of old HSCs. FACS analysis of HSCs from young and old mice indicated that the population of HSCs with higher CD150 levels significantly increases with aging (**Fig. 3b**), supporting the use of CD150 as an indicator of aging heterogeneity in old HSCs.

**Fig. 3.**
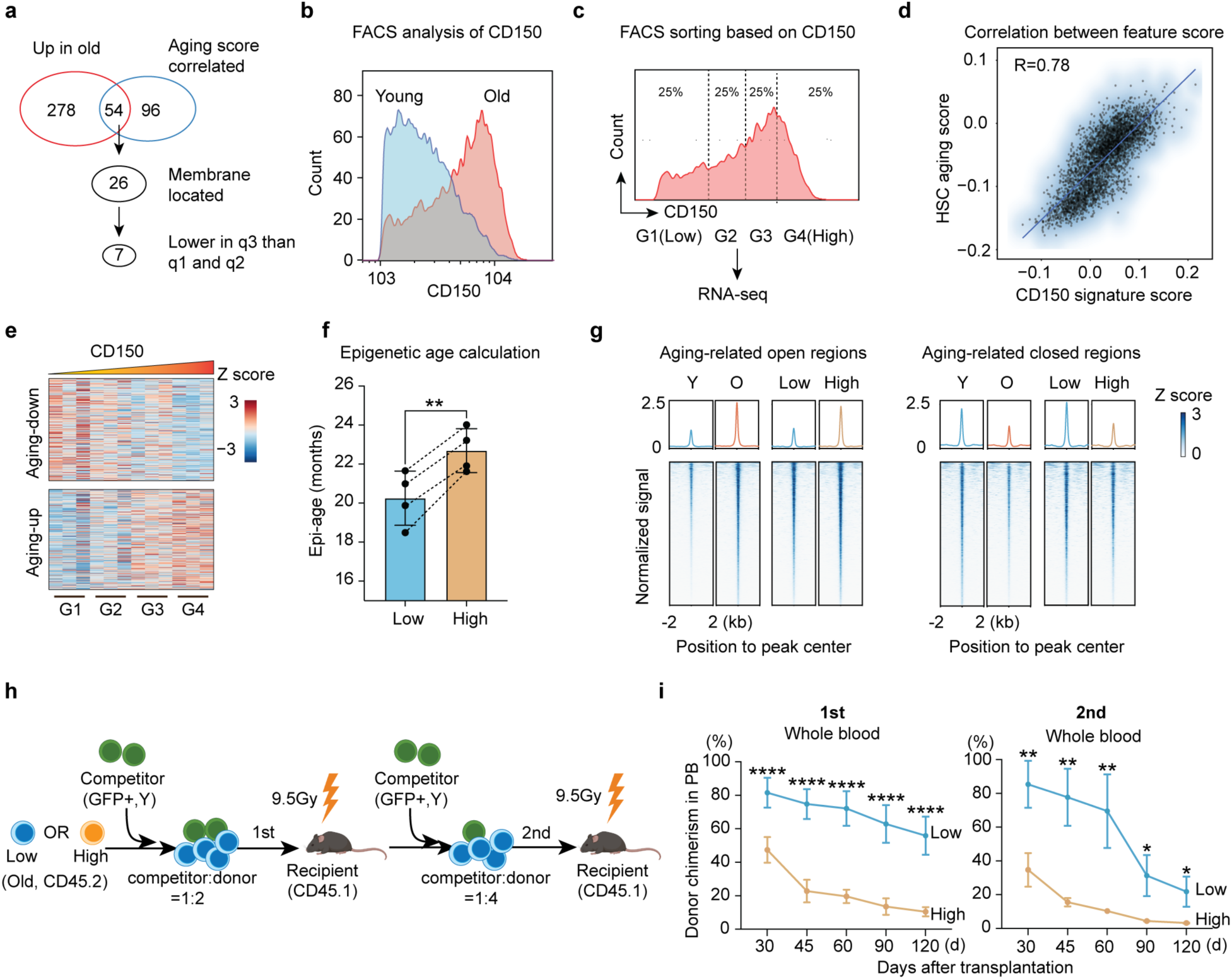
Identification of CD150 as a HSC aging heterogeneity marker. **a** Workflow for identifying heterogeneity marker genes in old HSCs. **b** FACS plot showing expression level of CD150 in young (3 months) and old (22-24 months) HSCs. **c** HSCs from old mice were separated into four subgroups (25% for each) based on their CD150 protein levels and subjected to bulk RNA-seq, *n* = 3. **d** Dot plot showing Pearson correlation between CD150 signature score and aging score based on scRNA-seq data, R=0.78. **e** Heatmap showing changes in expression of aging-related genes with ascending level of CD150 based on bulk RNA-seq in (C). **f** Bar plot showing the epigenetic age of CD150^low^ and CD150^high^ HSCs from 22-24 months old mice, *n* = 4, paired t test, the HSC subsets from the same mice were paired with dash line. **g** Line plot and heatmap showing ATAC-seq signal difference between CD150^low^ and CD150^high^ HSCs in aging-related open and closed regions, respectively. **h** Diagram of competitive transplantation to evaluate the repopulation capacity of CD150^low^ and CD150^high^ HSCs from old mice. **i** Whole blood chimerism of CD150^low^ and CD150^high^ HSCs from 22-24 months old donor mice at different time after the 1^st^ and 2^nd^ transplantation. *n* = 6 for the first and *n* = 3 for the second transplantation. Mean ± SD, student t test, \**P*<0.05, ** *P*<0.01, **** *P*<0.0001. The graphic of the mouse in **h** was created with BioRender.

To further confirm that the CD150 level can serve as a marker of aging heterogeneity of old HSCs, we separated old HSCs into 4 groups (G1 to G4) by FACS based on CD150 levels and profiled their transcriptomes (**Fig. 3c**). We then calculated the expression correlation between CD150 and each gene and identified 131 and 103 genes that exhibit strong positive or negative correlation with CD150 in aged HSCs, respectively (**Supplementary information, Fig. S5c; Supplementary information, Table S5**). Notably, the genes that positively correlate with CD150 include many known HSC aging markers, including *Sbspon, Ehd3, Clu, Selp, Jam2* and *Enpp5*. In contrast, the negatively correlated genes include genes that are highly expressed in young HSCs, such as *Serpinb1a, Usp6nl, Lrr1* and *Kif15* (**Supplementary information, Fig. S5d**). These results suggest that CD150 level can be a potential indicator of HSC transcriptome age.

Using the 131 genes that positively correlate with the CD150 level (referred to as CD150 feature genes), we calculated the CD150 feature score and found that q3 HSCs have a significantly lower CD150 feature score than q1 and q2 HSCs (**Supplementary information, Fig. S5e, f**). Notably, a strong correlation between the CD150 feature score and aging score was observed (**Fig. 3d**). This indicates that the aging heterogeneity of old HSCs can be well reflected by CD150 feature genes. Consistently, HSC aging-related up- and down-regulated genes in the 4 groups of HSCs from G1 to G4 exhibit a trend that is similar to the aging process (**Fig. 3e**), confirming the positive correlation between CD150 level and HSC aging status in old mice.

### CD150^low^ HSCs from old mice have younger epigenome and superior functions

Epigenetic changes are believed to be one of the drivers of aging.^47–49^ To further determine whether CD150 level can reflect epigenetic aging in old HSCs, we first compared their epigenetic age and found that CD150^low^ HSCs have a lower epigenetic age than that of CD150^high^ HSCs despite that they are from the same old mice (**Fig. 3f**). Given that stem cell aging is accompanied by chromatin accessibility changes,^50–52^ we compared the chromatin accessibility landscapes of old CD150^low^ and CD150^high^ HSCs, as well as young and old HSCs by performing ATAC-seq analyses. We found that 4,694 peaks displayed increased chromatin accessibility and 3,842 peaks showed decreased accessibility in aged HSCs relative to young HSCs, while 3,066 open and 1,966 closed differential peaks were identified between old CD150^low^ and CD150^high^ HSCs (**Supplementary information, Fig. S5g, h**). Interestingly, a similar change between CD150^low^ and CD150^high^ old HSCs was also observed around the aging-related ATAC-seq peaks (**Fig. 3g**). These results indicate that CD150 can serve as a marker reflecting both transcriptional and epigenetic aging in aged HSCs.

When we further check the genes close to the differential accessible peaks, we found several HSC aging-related genes, including *Clu*, *Jam2*, *Mt2*, *Nupr1*, *Selp*, *Slamf1*(CD150), and *Vwf* are located near the aging-related open peaks, and these genes also harbor the open peaks when comparing old CD150^high^ and CD150^low^ HSCs (**Supplementary information, Fig. S5g, h**). Consistently, the genes located around closed differential accessible peaks in both comparisons include many highly expressed genes in young HSCs, such as *Cd52*, *Cdc6*, *Haao* and *Nkg7*, demonstrating the corresponding chromatin accessibility changes underlying transcriptome alteration.

In addition, GO analysis of the genes related to the differential accessible peaks revealed enrichment of ‘cell adhesion’ and ‘potassium transport’ in the open peaks in both comparisons. Previous studies have demonstrated a correlation between ‘cell adhesion’ with HSCs functions and aging, ^46, 53, 54^ further supporting the potential functional defects in old and CD150^high^ HSCs compared to their counterparts.

To gain further insight into the aging-related open and closed ATAC-seq peaks, we performed motif enrichment analysis. We found that the aging-related open peaks are enriched for binding sites of the AP-1 family transcription factors (TFs), while the aging-related closed peaks are enriched for the ETS family TFs, including SPIB and PU.1 (**Supplementary information, Fig. S5i**). Importantly, similar TFs were identified when comparing CD150^low^ and CD150^high^ old HSCs (**Supplementary information, Fig. S5j**), further supporting a relatively younger chromatin in old CD150^low^ HSCs compared to the CD150^high^ HSCs. AP-1 family TFs have been reported as pioneer factors in aging,^55, 56^ while PU.1 and SPIB play important roles in regulating hematopoiesis process,^57, 58^ further highlighting the altered chromatin status and potentially impaired functions in the old CD150^high^ HSCs.

Previous studies have shown that increased DNA damage and decreased percentage of cells in proliferation are associated with functional decline of old HSCs.^27, 59, 60^ Interestingly, we observed an increase of double stranded breaks (indicated by γH2AX) as well as a decrease in the proportion of cells in proliferation in CD150^high^ compared to CD150^low^ old HSCs (**Supplementary information, Fig. S5k, l**), indicating CD150^low^ HSCs may have better functions than CD150^high^ HSCs in old mice.

To directly compare the functions between old CD150^low^ and CD150^high^ HSCs, we performed serial competitive transplantation (**Fig. 3h**). We found that CD150^low^ old HSCs have significantly better engraftment capacity compared to their CD150^high^ counterparts in both the first and second transplantation (**Fig. 3i; Supplementary information, Fig. S6a, b**). These results demonstrate that CD150 level not only marks transcriptome and epigenome heterogeneity, but also reflects functional heterogeneity of old HSCs.

Despite old CD150^low^ HSCs being molecularly and functionally younger than CD150^high^ HSCs, the extent to which they are similar to young HSCs remains unclear. To this end, we conducted molecular and functional comparisons between young HSCs and old CD150^low^ HSCs. The transcriptome comparison showed that, although old CD150^low^ HSCs are younger than CD150^high^ HSCs, they are still different from young HSCs in the expression of aging-related genes (**Supplementary information, Fig. S6c**). In terms of chromatin accessibility, old CD150^low^ HSCs are similar to young HSCs in aging-related closed regions but have higher signals in aging-related open regions (**Supplementary information, Fig. S6d**). These results demonstrate that old CD150^low^ HSCs are younger than old CD150^high^ HSCs, but still older than young HSCs at the molecular level. As for the direct functional comparison, based on the competitive transplantation experiment (**Fig. 3h**), we found that old CD150^low^ HSCs retain about 72.9% functionality relative to that of the young HSCs in terms of their repopulation capability, while the old CD150^high^ HSCs only retain about 5.9% (**Supplementary information, Fig. S6e**). Collectively, these results indicate that although old CD150^low^ HSCs are not as youthful as young HSCs at the molecular and functional levels, they retain most of the functionality of young HSCs.

Although young HSCs exhibit a homogeneous transcriptome (**Fig. 2d**) and aging score (**Fig. 2h**), their CD150 levels are variable (**Fig. 3b**). This raised the question whether CD150^low^ HSCs are also functionally superior to CD150^high^ HSCs in young mice. To address this question, we checked the cell cycle and performed similar HSC transplantation experiments using young HSCs (**Supplementary information, Fig. S7a, b**). Consistent with their transcriptome and aging score homogeneity, no significant functional difference was observed between CD150^low^ and CD150^high^ young HSCs for cell cycle analysis or peripheral blood (PB) chimerism during the period of 120 days after transplantation (**Supplementary information, Fig. S7a-d**). However, we noted a myeloid bias and significantly higher contribution to myeloid cells in young CD150^high^ HSCs, along with an upward trend in PB chimerism when compared to CD150^low^ young HSCs. These results indicate that CD150^high^ HSCs from young mice have comparable or even better functions compared to CD150^low^ HSCs, which is in contrast to those in old mice. Furthermore, these findings also suggest that myeloid-biased differentiation is not linked to aging status or decreased functionality in young HSCs.

### Old CD150^high^ HSCs are defective in differentiation, but not in self-renewal or activation

We next attempted to dissect the mechanism underlying the differential repopulation capacity of CD150^low^ and CD150^high^ HSCs in old mice. Considering that self-renewal and differentiation are the two major features of stem cells, we compared both features after their transplantation. We first determined the differentiation dynamics of old HSCs at different time points after transplantation (**Supplementary information, Fig. S8a**). We found that transplanted old HSCs start to differentiate on day 7, and the progenitor subpopulation becomes apparent on day 10 (**Supplementary information, Fig. S8b**). Based on this, we separately transplanted old CD150^low^ and CD150^high^ HSCs and analyzed the donor derived hematopoietic stem and progenitor cells (HSPCs) in bone marrow on days 7 and 14 after transplantation (**Fig. 4a**). Notably, we found that CD150^high^ HSCs showed a significantly lower ratio in short term HSCs (ST-HSCs) and multipotent progenitors (MPPs), but a higher ratio in LT-HSCs compared to CD150^low^ HSCs (**Fig. 4b; Supplementary information, Fig. S8c**), indicating that old CD150^high^ HSCs might have differentiation and/or activation defects.

**Fig. 4.**
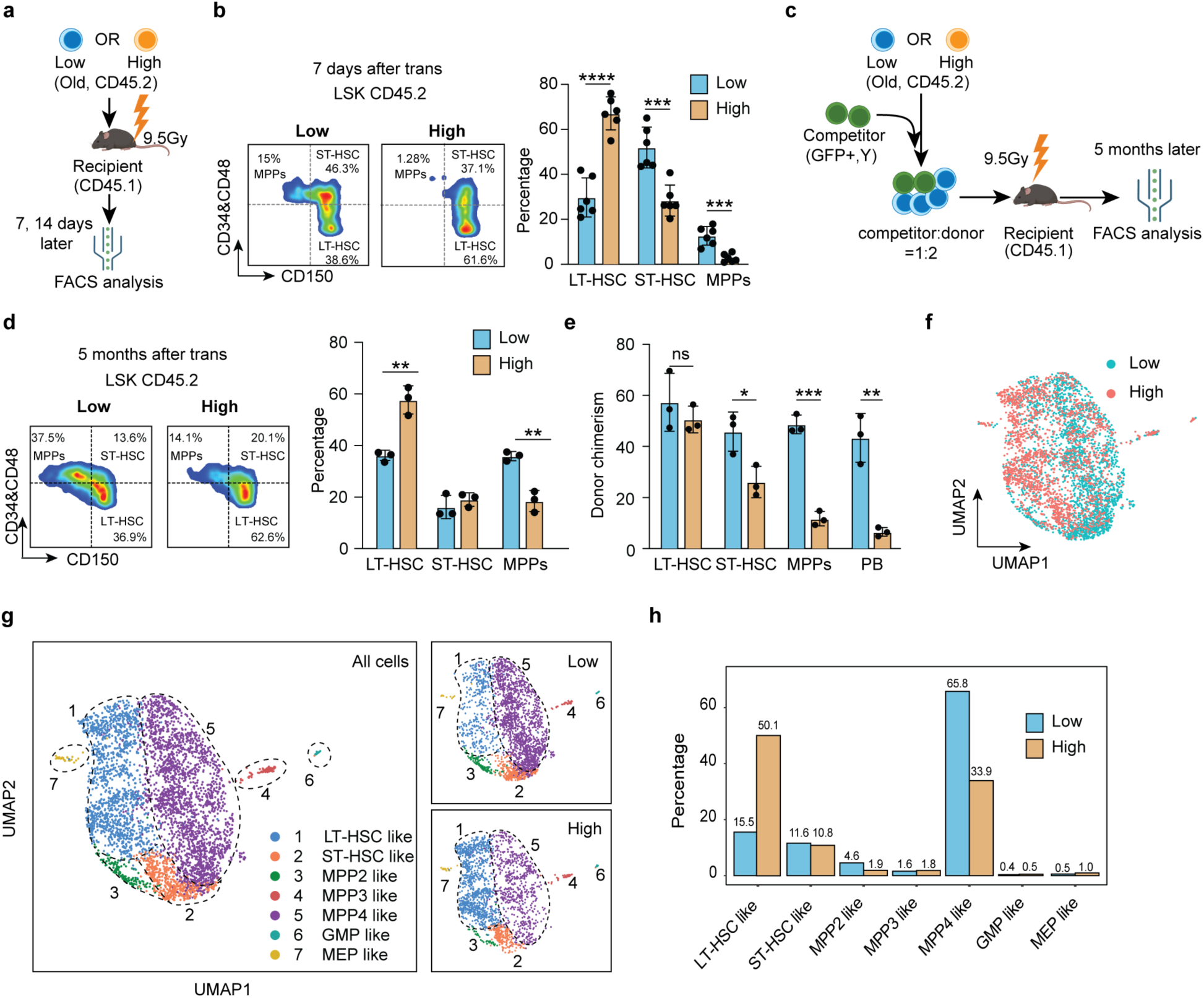
Differentiation, but not self-renewal, is a major defect of old CD150^high^ HSCs. **a** Diagram illustration of the transplantation experiment comparing old CD150^low^ and CD150^high^ HSCs. Donor HSCs derived HSPCs from the bone marrow were analyzed on days 7 and 14 after transplantation. **b** Representative FACS analysis of donor HSCs derived HSPCs (left) and quantification of different cell populations 7 days after transplantation (right). LT-HSC (CD45.2 LSK, CD34^-^CD48^-^CD150^+^), ST-HSC (CD45.2 LSK, CD34^+^/CD48^+^CD150^+^) and MPPs (CD45.2 LSK, CD34^+^/CD48^+^CD150^-^) were analyzed, *n* = 3. **c** Diagram illustration of the competitive transplantation for evaluating the long-term differentiation of CD150^high^ HSCs. Both peripheral blood and bone marrow were analyzed five months after transplantation. **d** Representative FACS analysis of donor HSC-derived HSPCs (left) and quantification of different cell populations (right) 5 months after transplantation, *n* = 3. **e** Bar graph showing the chimerism of donor HSCs in LT-HSCs, ST-HSCs, MPPs and PB 5 months after transplantation, *n* = 3. **f** UMAP plot showing cell distribution of HSPCs derived from CD150^low^ and CD150^high^ HSCs in scRNA-seq 14 days after transplantation. **g** UMAP presentation of cell types predicted based on HSPC marker gene expression (left). In total, 7 different cell types were identified in HSPCs. **h** Bar graph showing relative abundance of predicted cell types in HSPCs from mice that received old CD150^low^ or CD150^high^ HSCs. Mean ± SD, student t test, \**P*<0.05, \*\**P*<0.01, \*\*\**P*<0.001. The graphic of the mouse in **a** and **c** were created with BioRender.

To distinguish between these two possibilities, we profiled transcriptomes of CD150^low^ and CD150^high^ HSCs 4 days after transplantation when the HSCs had yet to initiate differentiation (**Supplementary information, Fig. S8d**). By comparing with freshly isolated HSCs, we identified 794 commonly up-regulated genes in both CD150^low^ and CD150^high^ HSCs after transplantation (**Supplementary information, Fig. S8e; Supplementary information, Table S6**). The commonly up-regulated genes are highly enriched in terms related to cellular division. Consistently, the G2M checkpoint related genes were similarly upregulated in both CD150^low^ and CD150^high^ HSCs upon transplantation (**Supplementary information, Fig. S8f**), these results indicate that old CD150^high^ HSCs can be effectively activated as that of CD150^low^ HSCs following transplantation.

We further compared the proliferation capacity of old CD150^low^ and CD150^high^ HSCs in culture and found that old CD150^low^ and CD150^high^ HSCs exhibited comparable proliferation rates (**Supplementary information, Fig. S8g-k**). Taken together, these results indicate that impaired repopulation capacity of old CD150^high^ HSCs is caused by defective differentiation, but not activation or proliferation.

To rule out a possible delay in differentiation of old CD150^high^ HSCs after transplantation, we analyzed the persistence of the differentiation and self-renewal capacity of old CD150^high^ HSCs over an extended period of 5 months. We co-transplanted old CD150^low^ or CD150^high^ HSCs (GFP-) with competitors (GFP+) and analyzed the differentiation and chimerism of donor HSCs 5 months later (**Fig. 4c**). We observed a consistent differentiation defect of old CD150^high^ HSCs when compared to their CD150^low^ counterparts by analyzing donor derived HSPCs in the bone marrow (**Fig. 4d**). Additionally, the donor chimerism of old CD150^high^ HSCs exhibits a clear trend of decline from LT-HSCs through ST-HSCs, MPPs, to PB, while the donor chimerism of old CD150^low^ HSCs remains relatively stable (**Fig. 4e**), supporting a long-term differentiation defect of old CD150^high^ HSCs.

Collectively, these results indicate a long-lasting impairment in differentiation, but not in activation or self-renewal of old CD150^high^ HSCs.

### Old CD150^high^ HSCs are defective in the LT-HSCs to ST-HSCs transition

To determine the specific stage when the differentiation defects of old CD150^high^ HSCs occur, we performed comparative scRNA-seq analysis of the HSPCs derived from old CD150^low^ and CD150^high^ HSCs 14 days after transplantation. We obtained 2,406 and 1,925 high quality HSPCs derived from transplanted CD150^low^ and CD150^high^ HSCs, respectively (**Fig. 4f**). Cell cycle phase analysis indicated that similar proportions of HSPCs derived from CD150^low^ and CD150^high^ HSCs were actively cycling (**Supplementary information, Fig. S9a**), consistent with our previous results (**Supplementary information, Fig. S8e, f**).

Based on the expression of the HSPC marker genes ^50, 61^ and the HSPC sorting markers (*Ly6a*, *Kit*, *Cd34*, *Cd48* and *Slamf1*), the 4,331 HSPCs can be divided into 7 subclusters that include LT-HSC-like, ST-HSC-like, MPP2-like, MPP3-like, MPP4-like, GMP-like and MEP-like cells (**Fig. 4g; Supplementary information, Fig. S9b**). When HSPCs derived from CD150^low^ and CD150^high^ HSCs were compared, we found that HSPCs derived from CD150^high^ HSCs were significantly enriched in LT-HSC-like cells (**Fig. 4g, h**), consistent with our FACS analysis (**Fig. 4b; Supplementary information, Fig. S8c**). Since the proportions of CD150^low^ and CD150^high^ HSCs derived ST-HSCs are comparable, our data indicate that the differentiation defects of CD150^high^ HSCs were mainly at the stages of LT-HSC to ST-HSC, and ST-HSC to MPP4 transitions. Given that MPP4 are the major progenitor cells of the lymphoid lineage, the substantial decrease in the MPP4 population might be a major cause of the biased differentiation of CD150^high^ HSCs.^30, 32^

Consistently, when we further performed pseudo time analysis to map the differentiation trajectory, we found that HSPCs derived from CD150^high^ HSCs were predominantly enriched at early stages and less enriched at later stages compared to HSPCs derived from old CD150^low^ HSCs (**Supplementary information, Fig. S9c**), supporting that the differentiation of CD150^high^ HSCs is impaired compared to their CD150^low^ counterparts.

To further analyze the transcriptome differences between populations derived from old CD150^low^ and CD150^high^ HSCs, we identified differentially expressed genes between them. Significant transcriptome differences were observed in LT-HSC-like, ST-HSC-like, and MPP4-like subpopulations (**Supplementary information, Fig. S9d-f**). Interestingly, some HSC aging marker genes, such as *Vwf*, *Nupr1*, *Ifitm3*, and *Ifitm1* are consistently upregulated in HSPCs derived from old CD150^high^ HSCs, indicating that upregulation of HSC aging genes can persist in downstream cells.

To further investigate the transcriptome difference, we performed Gene Ontology (GO) analysis. Upregulated genes in cells derived from CD150^low^ HSCs are consistently enriched for ‘translation-related pathways’ across cell types, while those from CD150^high^ HSCs are enriched for ‘protein folding response-related pathways’ (**Supplementary information, Fig. S9g**). Increased translation in HSCs is associated with differentiation activities,^62–64^ indicating more active differentiation in old CD150^low^ HSCs. Protein folding response is linked to cellular stress, a hallmarks of HSC aging,^65, 66^ consistent with the impaired functions of old CD150^high^ HSCs. The consistently enriched pathways among cell types indicate that transcriptomic defects in HSCs may be passed to downstream progenitors.

To gain further support for this notion, we performed Gene Set Enrichment Analysis (GSEA) comparing HSPCs from old CD150^low^ and CD150^high^ HSCs. Several pathways are consistently enriched in cells derived from old CD150^high^ HSCs, including IFN-α response, mTOR signaling, unfolded protein response, and glycolysis (**Supplementary information, Fig. S9h**). Some of these pathways have been linked to stem cell aging and their differentiation activities,^66–69^ further supporting that transcriptome defects in HSCs can be passed on to downstream cells.

Collectively, comparative bone marrow scRNA-seq analysis of transplanted old HSCs indicated that the differentiation defects of old CD150^high^ HSCs is caused by a block of LT-HSCs to ST-HSCs transition and the transcriptome defects of HSCs can be passed on to downstream progenitor cells.

### The “younger” subset of old HSCs can alleviate aging phenotypes and extend lifespan

Transplantation of young HSCs as an anti-aging therapy is impractical for humans due to immune incompatibility among different individuals. Given this challenge, the demonstration that HSCs in old mice have molecularly and functionally “younger” CD150^low^ HSCs prompted us to ask whether this subset of HSCs can help to attenuate aging phenotypes of old mice. To this end, we transplanted CD150^low^, whole HSCs (un-selected), or CD150^high^ HSCs from old mice into 13-month-old recipients and analyzed aging phenotypes 5 months later (**Fig. 5a**). Compared to mice that received CD150^high^ HSCs, mice transplanted with CD150^low^ HSCs showed higher percentage of B cells, and lower percentage of T cells and myeloid cells (**Fig. 5b**), as well as increased blood cell numbers, including lymphoid cells, neutrophils, eosinophils and red blood cells (**Fig. 5c**). Notably, a significant increase in naïve T cell ratio and a corresponding decrease in the Tcm and Tem cell ratios were observed when comparing mice that received old CD150^low^ HSCs to those received old CD150^high^ HSCs (**Fig. 5d; Supplementary information, Fig. S10a**). These results suggest that transplantation of old CD150^low^ HSCs can improve both hematopoiesis and immune parameters in old recipient mice.

**Fig. 5.**
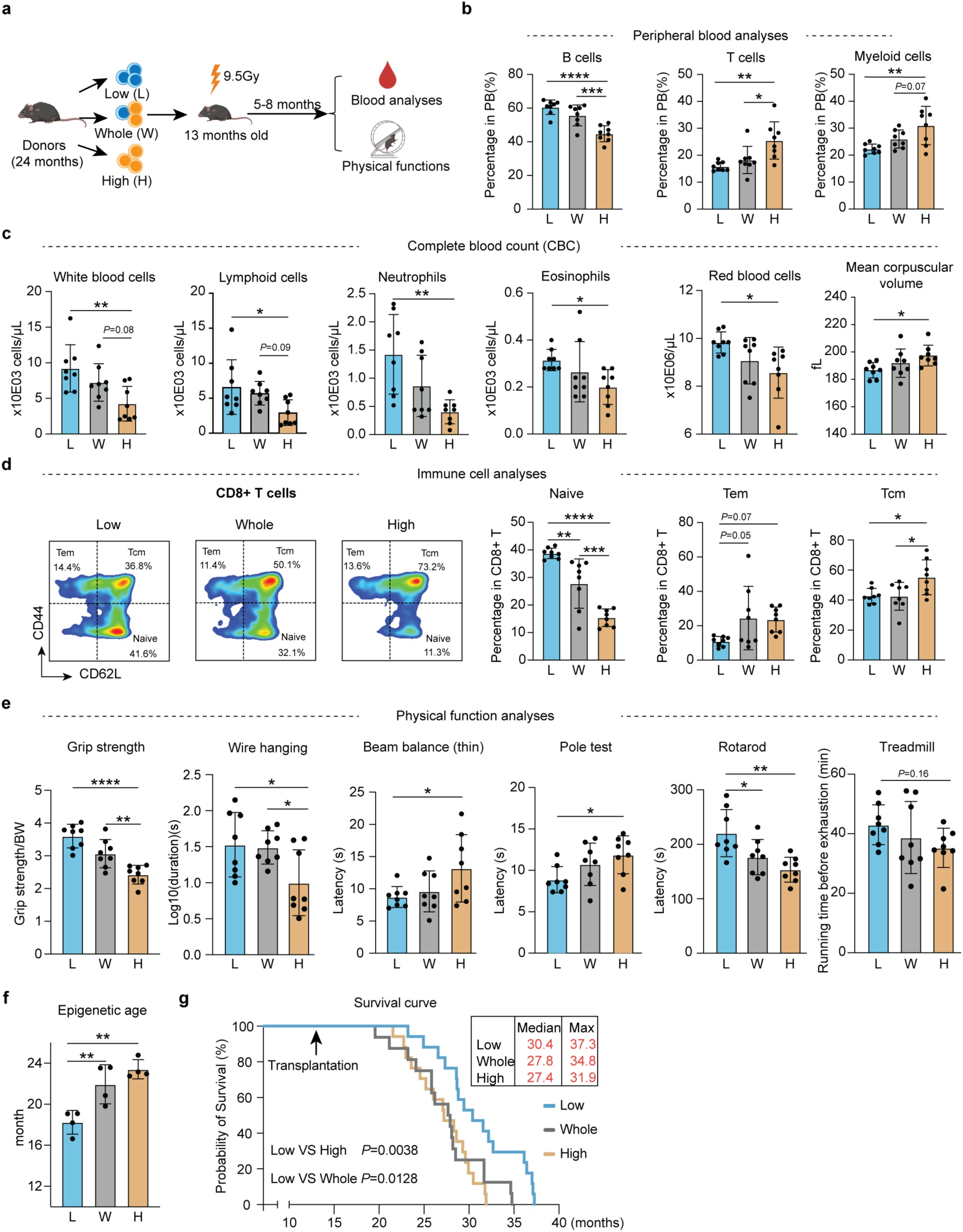
Transplantation of “younger” subset old HSCs alleviates aging phenotypes of aged mice. **a** Diagram showing the experiment design of individual transplantation of 2,000 old CD150^low^ (25% lowest), whole-HSCs (un-selected) and CD150^high^ (25% highest) HSCs into middle-aged recipients (13-month-old), *n* = 8. Five months after transplantation, a series of hematopoietic and physical tests were performed. **b** Bar graph showing the percentage of B, T and myeloid cells in the PB of recipient mice of the three groups, *n* = 8. **c** Bar graph showing absolute number of blood cells in PB of recipient mice from different transplanted groups, *n* = 8. **d** Representative FACS analysis of naïve T cells, central memory T cells (Tcm) and effector T cells (Tem) ratio in CD8 positive T cells and their quantification in recipient mice of the three groups, *n* = 8. **e** Physical tests of recipient mice that received CD150^low^, whole-HSCs and CD150^high^ HSCs. Muscle strength, motor coordination and endurance were assessed, *n* = 8. **f** Bar graph showing the epigenetic age of recipient mice from different groups using blood samples, *n* = 4. **g** Survival curve showing the lifespan difference among mice that received old CD150^low^, whole-HSCs and CD150^high^ HSCs, each mouse received 2,000 HSCs. Log-rank (Mantel-Cox) test, *n* = 17. For (**b-f**), Mean ± SD, one-way ANOVA, \**P*<0.05, \*\**P*<0.01, \*\*\**P*<0.001, \*\*\*\**P*<0.0001. The graphic of the mouse in **a** was created with BioRender.

To determine whether improved hematopoiesis and immune functions can improve physical performance, we performed a series of tests that measure muscle strength (grip strength, wire hanging), coordination (beam balance, pole test), and endurance (rotarod and treadmill). Consistently, we found a general trend of degrading physical performance in the order of CD150^low^, whole-HSCs, and CD150^high^ HSCs (**Fig. 5e**) without significant differences in their body weight (**Supplementary information, Fig. S10b**). Consistent with improved physical performance, we found that old mice that received CD150^low^ HSCs display a lower epigenetic age in blood (**Fig. 5f**). Importantly, aged recipient mice transplanted with old CD150^low^ HSCs exhibited a substantial lifespan extension, achieving a 9.4% and 11.5% increase in median lifespan, and a 7.2% and 16.9% increase in maximum lifespan compared to counterparts transplanted with whole-HSCs or CD150^high^ HSCs (**Fig. 5g**).

Taken together, our results indicate that not only transplantation of young HSCs, but the “younger” subset of old HSCs can attenuate aging phenotypes of old mice. Considering that the functionally defective CD150^high^ HSCs increase with aging, our study raises the possibility that aging-alleviating effects might be achieved by reducing or removing the functionally defective CD150^high^ HSCs from old mice.

### Reducing the dysfunctional HSCs can alleviate aging phenotypes in old recipient mice

To assess whether reducing defective HSCs in old mice can enhance hematopoiesis and mitigate aging-related phenotypes, we mixed 500 old CD150^low^ HSCs with 2,500 (1:5 ratio), 1,000 (1:2 ratio) or 0 (1:0 ratio) CD150^high^ old HSCs and co-transplanted to irradiated middle-aged (13 months) mice to mimic whole-HSC, partial removal, and complete removal of CD150^high^ HSCs, respectively. Five months after the transplantation, we assessed the hematopoietic system and physical functions of recipient mice to compare the aging status of the three groups (**Fig. 6a**). The 1:5 ratio serves as a control, which is determined based on the percentage of q3 to the total HSC in old mice based on scRNA-seq (**Fig. 2d**).

**Fig. 6.**
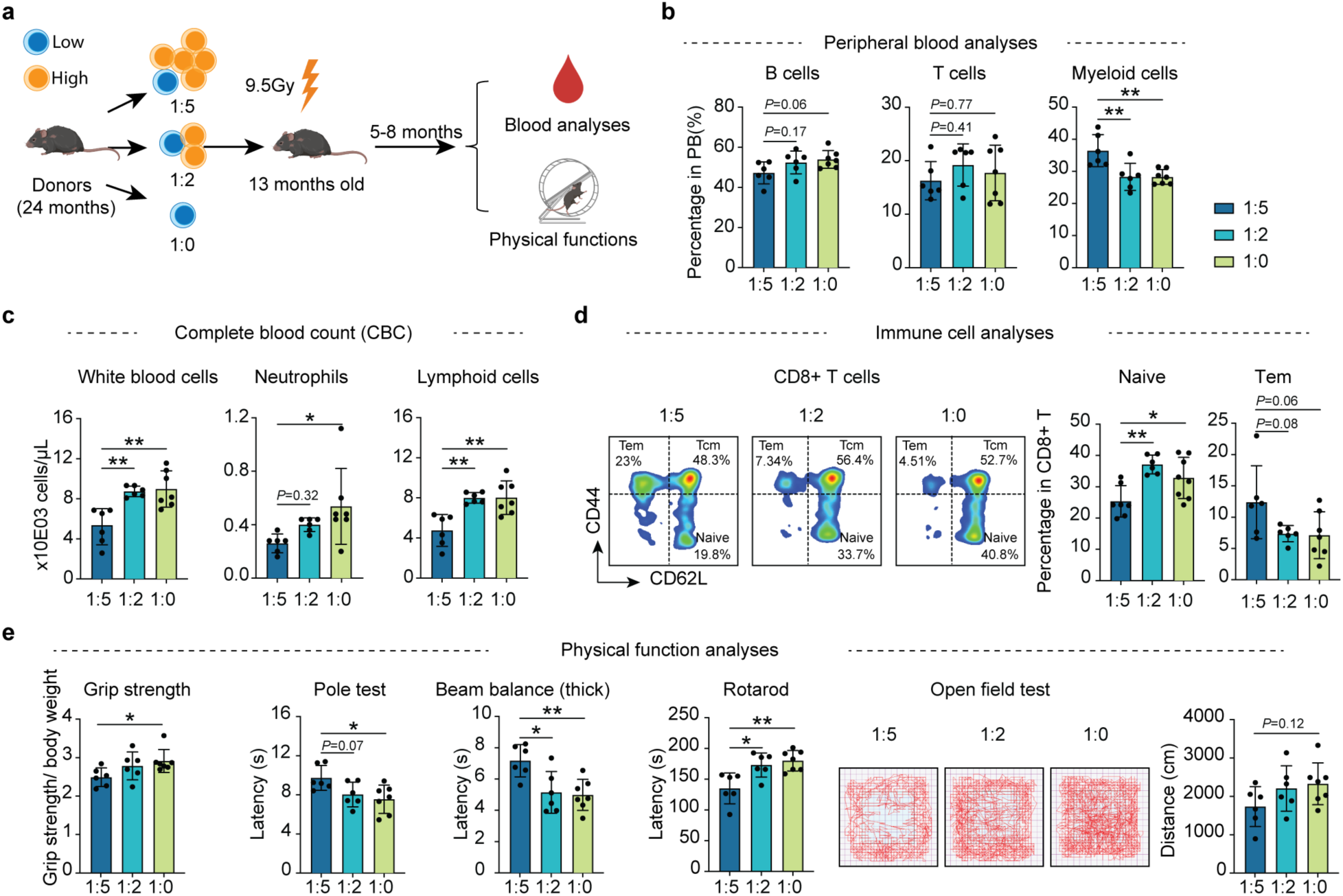
Reducing dysfunctional HSCs ameliorates aging phenotypes in old mice after transplantation. **a** Diagram showing the experiment design. Individual transplantation of 500 old CD150^low^ HSCs with: 2,500 CD150^high^ HSCs (1:5 group); 1,000 CD150^high^ HSCs (1:2 group) and 0 CD150^high^ HSCs (1:0 group) into middle-aged recipients (13-months-old). Five months after transplantation, a series hematopoietic and physical tests were performed. **b** Bar graph showing the percentage of B, T and myeloid cells in the PB of recipient mice of the different groups. **c** Bar graph showing the blood cell numbers in PB of recipient mice from different transplanted groups. **d** Representative FACS analysis of naïve T cells and effector T cells (Tem) ratio in CD8 positive T cells and their quantification in recipient mice of different groups. **e** Bar plot showing the results of physical functional tests of recipient mice of different groups. Muscle strength, motor coordination, endurance and locomotor activity were assessed. *n* = 6 for 1:5 and 1:2 group, *n* = 7 for 1:0 group, Mean ± SD, one-way ANOVA, \**P*<0.05, \*\**P*<0.01. The graphic of the mouse and equipment in **a** were created with BioRender.

Peripheral blood analyses indicated that compared to the 1:5 group, the 1:2 and 1:0 groups have increased percentage of B cells and decreased percentage of myeloid cells (**Fig. 6b**). Complete blood counting also showed higher number of whole white blood cells, neutrophils, and lymphoid cells in the 1:0 and 1:2 groups compared to the 1:5 group (**Fig. 6c**). Importantly, increased naïve CD4^+^ and CD8^+^ T cells, as well as decreased effective memory T cells were also detected in the 1:2 and 1:0 groups compared to the 1:5 group (**Fig. 6d; Supplementary information, Fig. S11a**), indicating the presence of the defective CD150^high^ HSCs has adverse effects on the “younger” functional CD150^low^ HSCs for hematopoiesis and immune function in old mice.

Next, we examined whether reducing the CD150^high^ HSCs ratio can improve physical functions of the recipient mice by performing a multitude of physical tests. The results indicate consistent improved physical functions in the 1:2 and 1:0 groups compared to the 1:5 group, including muscle strength (grip strength), motor coordination (pole test, beam balance, and rotarod), and locomotor activity (open field test), with no notable differences in body weight (**Fig. 6e; Supplementary information, Fig. S11b**). These findings not only confirm the adverse effect of the defective CD150^high^ HSCs in aged mice, but also suggest that their reduction in old mice can ameliorate aging-related phenotypes.

## Discussion

By transplantation, we demonstrated a key role of HSCs in systematic aging. Integrating scRNA-seq and functional evaluation, we identified a “younger” subset of HSCs in old mice that is marked by low level of CD150. Comparative analysis demonstrated that CD150^high^ HSCs from old mice are defective in LT-HSC to ST-HSC differentiation compared to the CD150^low^ HSCs. Importantly, transplantation of the “younger” CD150^low^ HSCs can alleviate aging phenotypes and extend the lifespan of old mice. Finally, we provide evidence demonstrating that the hematopoietic system and physical performance of old recipient mice can be improved by reducing the ratio of functionally defective CD150^high^ HSCs, raising the possibility that rejuvenation can be achieved by removing the dysfunctional HSCs in the elderly.

Originally characterized as a cell surface marker for LT-HSCs,^70^ CD150 has also been used to reflect differentiation bias of HSCs in young and old mouse.^30, 32^ Additionally, CD150^low^ HSCs from aged mice have been shown to retain normal lymphoid differentiation capacity when removed from the aged microenvironment.^34^ These previous results suggest a potential link between CD150 levels and HSC function during aging and are in line with our results, which further support CD150 as a marker of HSC aging heterogeneity. In addition to CD150, we identified other cell surface proteins—such as *Ramp2*, *Ehd3*, *Efna1*, *Gpr183*, *Jam2*, and *Tm4sf1*—as potential markers of aging heterogeneity in old HSCs. Since all these markers were identified using the same criteria and they exhibit strong positive correlations with each other in expression (data not shown), we believe they would segregate old HSCs into very similar populations. We chose to use CD150 over the others due to the availability of antibodies and their performance on old HSCs sorting (data not shown). Interestingly, during the preparation of this manuscript, Kim *et al*. reported that *Clca3a1* can also serve as a marker to capture the HSC heterogeneity in old mice. They found that Clca3a1^high^ HSCs from aged mice have a decreased repopulation capacity in primary transplantation and exhibit a myeloid-biased differentiation compared to Clca3a1^low^ HSCs.^31^ However, this functional difference between Clca3a1^high^ and Clca3a1^low^ old HSCs was not maintained in the second transplantation. In contrast, the functional difference between CD150^high^ and CD150^low^ old HSCs persist in the second transplantation, indicating that CD150 is a more robust indicator of the functional heterogeneity of aged HSCs.

Since the discovery of the rejuvenating effects of young blood through parabiosis,^7^ great efforts have been spent on searching for the rejuvenation factors from young blood. Although modulating several candidate factors, such as CDC42, SELP, CCL5, PHF6, IGF1, PF4 and the Yamanaka factors (Oct4, Sox2, Klf4, c-Myc) have been reported to have rejuvenating effects during the past years,^23, 46, 71–77^ their rejuvenating capacities are limited and the mechanisms underlying their rejuvenating effects are not clear. In some cases, such as GDF11, conflicting results were also reported.^9, 10, 78, 79^ Thus far, no universally accepted rejuvenating factors have been identified. Our findings reveal that old HSCs exhibit molecular and functional heterogeneity, comprising “younger” and dysfunctional “older” subsets. Moreover, reducing the proportion of dysfunctional HSCs ameliorates aging phenotypes in aged mice after transplantation, suggesting that removal of functionally defective HSCs from old could be a promising approach for alleviating aging phenotypes. Consistent with this notion, a very recent study showed that depletion of the myeloid-biased HSCs (marked by NEO1) by a specific antibody cocktail could restore immune functions in old mice,^80^ although whether such approach can lead to a systemic body rejuvenation remains unknown. Nevertheless, our study and this recent report have set the stage for achieving rejuvenation in the elderly by targeted removal of dysfunctional HSCs.

## Material and methods

### Mice

All experiments were conducted in accordance with the National Institute of Health Guide for Care and Use of Laboratory Animals and approved by the Institutional Animal Care and Use Committee (IACUC) of Boston Children’s Hospital and Harvard Medical School. For bulk and single-cell RNA-seq, 2-3 months young and 22-24 months old male C57BL/6 mice were used (Jackson Lab #00664). For HSC transplantation, 2-3 months old male young (Jackson Lab #00664) or 2-3 months old GFP+ male young (Jackson Lab #006567) mice were used. For competitive transplantation, competitor HSCs were collected from 3-4 months GFP+ male young (Jackson Lab #006567) mice. Except specially mentioned, the recipient mice were B6 CD45.1 male mice (Jackson Lab #002014), and the helper cells were also collected from B6 CD45.1 male mice. For old recipient mice, 13 months old wild type male mice were used and the helper cells were from age-matched wild type male mice.

### LT-HSCs sorting and transplantation

Bone marrow cells were collected by crushing tibias, femurs, pelvic bone and spine. To improve the sorting efficiency, collected cells were further enriched by removing lineage positive cells (CD4, CD8, Gr-1, CD11b, CD5, B220 and Ter119) with selection beads (STEMCELL catalog #19856). The enriched cells were further stained with antibodies against mouse c-Kit, Sca-1, CD48, CD34, CD150. The LT-HSCs were defined as Lin^-^Sca-1^+^c-Kit^+^CD48^-^CD34^-^CD150^+^. For transplantation experiments, LT-HSCs were sorted with FACS sorter (SONY SH800), mixed with 3×10e5 Sca-1 depleted helper cells and transplanted into lethally irradiated (9.5 Gy) recipient mice. For the HSPCs sorting for 10X Genomics single-cell library preparation, we first transplanted old CD150^low^ or CD150^high^ HSCs (CD45.2) into irradiated young recipient (CD45.1) mice. Fourteen days later, bone marrow cells were collected, and the lineage-positive cells were removed using selection beads. The remaining cells were then stained with antibodies targeting CD45.1, CD45.2, Sca-1, and c-Kit, and CD45.2 LSK cells were sorted for 10x Genomics library preparation.

### Cell cycle and DNA damage marker **γ**H2AX analysis of HSCs with FACS

The analysis of cell cycle status of HSCs was performed as pervious described^81^. Briefly, bone marrow cells were prepared, enriched, and stained with HSC marker antibodies as mentioned before. Then the enriched bone marrow cells were fixed and permeabilized with BD Intra Kit (#641776). After washing, fixed HSCs were stained with monoclonal antibody against mouse KI67 (eBioscience, #11-5698-80) and Hoechst33342 (Thermo Scientific, #R37165) or with antibody against DNA damage marker γH2AX (Cell Signaling Technology, #9719S). The stained cells were analyzed with FACS after washing.

### Bulk RNA-seq library preparation

Bulk-RNA sequencing library is constructed based on Smart-seq2.^82^ SMART-Seq v4 Ultra Low Input RNA Kit for Sequencing (Takara catalog #634890) was used and experiments were performed according to manufacturer’s instructions.

### scRNA-seq library preparation

Single-cell RNA seq library is prepared with Chromium Next GEM Single Cell 3’ Reagent Kits V3.1 (Dual Index) from 10X Genomics according to manufacturer’s instruction and previously described method.^83^

### Peripheral blood analysis via FACS

For PB analysis, up to 30 µL vein blood was collected from retro orbital sinus or tail tips into EDTA-coated tubes. Red blood cells were first removed by red blood cell lysis buffer (ebioscience catalog #00-4333-57). The remaining white blood cells were stained with mixed monoclonal conjugated antibodies (CD45.1, CD45.2, CD3, B220, Gr-1 and CD11b). After incubation in 4°C and darkness for 30 min, PB were analyzed by FACS (BD FICR canto-II).

### HSC culture in vitro

Sorted LT-HSCs were cultured in StemSpan™ SFEM II (STEM CELL catalog #09605) with 10 ng/mL SCF (PeproTech, catalog #AF-250-03), 100 ng/mL TPO (PeproTech, catalog #315-14), 25 ng/mL Flt3L (PeproTech, catalog #250-31L) and 25 ng/mL IL3 (PeproTech, catalog # 213-13). Six days after culture, the number of HSCs were quantified by FACS (BD FICR canto-II).

For the proliferation curve, single HSC was sorted and cultured. The cell number was counted and recorded every day.

### Complete blood counting analysis of recipient mice

For whole mouse blood analysis, 75 µL vein blood was collected from retro orbital sinus and mixed with 225 µL 5mM EDTA. Freshly collected whole blood was analyzed by Hematology system (ADVIA 120) within 2 hours after collection.

### Naïve T, Tcm and Tem analysis

We analyzed the ratio of Naïve T cells, Tcm and Tem based on a pervious report.^84^ Briefly, PB was collected and red blood cells were removed by red blood cell lysis buffer (ebioscience catalog #00-4333-57). The remaining cells were stained with antibodies against CD45.1, CD45.2, CD4, CD8, CD44 and CD62L. T cell subtypes were defined as follows: naïve (CD62L^high^ CD44^low^), Tcm (CD62L^high^ CD44^high^) and Tem (CD62L^low^ CD44^high^).

### Physical function tests of mouse

Grip strength was tested by placing mice on a grid strength meter (Bioseb, Model GT3) so only their forepaws grasped the grid. Their tails were pulled three times, then they rested for at least 1 minute. We averaged the top three maximum grip strengths and normalized for body weight.

Beam balance test was performed with homemade equipment: a 1-meter smooth wood strip elevated 50 cm. Mice were trained on a thicker pole (28 mm diameter) until they could cross without dropping. The next day, they were tested three times on the thicker or thinner pole (17 mm diameter). Latent time to cross was recorded.

For pole test, mice were placed at the top of a 50 cm grooved metal pole with a diameter of 1□ cm with the mice head pointing down. The time from initial placement on the top of the pole to the time the mouse reached the base of the pole (forelimb contact with platform) was recorded with a stopwatch. Following a 30-min rest in their home cage, the trial was repeated another 2 times. The average time is calculated as the latent time to go down for each mouse.

The rotarod test assessed motor coordination and balance. Mice were placed on an accelerating Rotarod (Ugo Basile Apparatus) starting at 4 rpm. On day one, mice trained at 4 rpm for 5 minutes, repeated twice. On test day, the rod accelerated from 4 to 40 rpm over 5 minutes. Time and speed were recorded when mice dropped or after two passive rotations. Each mouse performed three tests over two days. Results were averaged over these tests.

For the wire hanging test, mice were placed on a metal grid with 1 cm mesh, elevated 50 cm above a cushioned surface with ∼3 cm bedding. Mice were put on the metal grid, which was slowly inverted so they hung upside down. The time until the mouse fell was recorded. Each mouse was tested three times with 30-minute intervals, and results were averaged.

For treadmill fatigue test, the protocol was adapted and modified from a previous publication.^85^ Briefly, mice were acclimated to the treadmill (Columbus Instruments) for 2 days before testing. The fatigue zone was set one mouse body-length from the shock grid. On day 1, mice explored the treadmill for 5 minutes with the shock grid off, then ran at 6 m/min for 5 minutes with the shock grid on (1.5 mA, 3 Hz). Speed increased by 2 m/min every 2 minutes, stopping when mice ran at 10 m/min for 5 minutes. On day 2, the same protocol was followed with a top speed of 12 m/min. On the test day, speed increased stepwise up to a maximum of 26 m/min. Fatigue was defined when mice remained in the fatigue zone for over 5 seconds on three occasions. Latency to fatigue was recorded for analysis.

For the fear conditioning test, we used contextual and cued fear conditioning (CCFC). On day one, mice underwent 2 minutes of acclimation, followed by a 30-second tone and a 2-second foot shock (0.5 mA), then a 2-minute intertrial interval; this sequence was repeated once more. The next day, mice were placed in the same context for 5 minutes without shock or tone (context test). On day three, mice were assessed in a modified context: 3 minutes without tone, then 3 minutes with tone (cued test). Video was captured using Noduls MediaRecorder 4.0, and freezing behavior was analyzed using Noldus EthoVision XT17.

For the Novel Y-Maze test, adapted from a previous study,^86^ mice were pre-acclimated in a separate holding room for 30 minutes. The procedure included a 3-minute habituation phase with one arm blocked, followed by a 3-minute test phase, separated by a strict 2-minute intertrial interval. Before the test phase, the obstruction was removed, the maze cleaned, and mice were placed back into the start arm for the test trial. Distance and time traveled were recorded and analyzed using Noldus EthoVision XT17.

Open field testing used a transparent enclosure with a base of 27.3□cm × 27.3 cm and walls 20.3□cm high; the central zone was half of the total area. Mice were acclimated for at least 20 minutes prior to testing. Each mouse was placed in the center to begin assessment. Movements were tracked for 20□ minutes using a tracking system (Med Associates, ENV-510), with data collected every 5 minutes. Analysis included total distance moved, average speed, active and inactive periods, and vertical events during the first 10 minutes.

### Reduced-representation bisulfite sequencing (RRBS) library preparation for HSCs

FACS sorted HSCs were lysed at 50°C for 3 hours and 75°C for 30 min in 5 μL lysis buffer (20 mM tris-EDTA, 20 mM KCl, 0.3% Triton X-100, and 1 mg/mL protease). The lysed cells were then subject to Msp I digestion at 37°C for 3 hours and 80°C for 20 min. Following enzymatic digestion, end-repair mix (1 μL Klenow frag exo (5 U/μL), 0.2 μL of 10×TangoBuf, and 0.8 μL of dNTP mix) was added and incubated at 37°C for 40 min and 75°C for 15 min. For adaptor ligation, the reaction was incubated at 16°C for 30 min, 4°C overnight, and 65°C for 20 min after adding the ligation mix (2.25 μL H_2_O, 1 μL T4 ligase (30 U/μL), 0.5 μL 100 mM ATP, and 0.75 μM methylated adaptor). Bisulfite conversion reaction was then performed by the EpiTect Bisulfite Kit (Qiagen catalog #59104) following the manufacturer’s instructions. Adapter-tagged BS-DNA was then amplified by 16 cycles using Kapa HiFi Uracil+ ReadyMix (Kapa Biosystems, catalog #KK2801) with NEBNext Multiplex Oligos for Illumina (New England BioLabs catalog # E7500L).

### ATAC-seq library construction

About 500 ∼ 1000 cells were sorted by FACS to 10 µL tagmentation buffer (33 mM Tris-Acetate, 66 mM K-Acetate, 10 mM Mg-Acetate, 16% Dimethylformamide, and 0.02% Digitonin). Then, 0.5 µL Tn5-Adapter complex (Diagenode) was added and incubated at 37°C for 30 min. The tagmentation reaction was stopped by adding 10 µL stop buffer (100 mM Tris-HCl pH8.0, 100 mM NaCl, 0.4% SDS, 40 µg/ mL Proteinase K) and the samples were incubated at 55°C overnight to release the fragments. SDS was then quenched by adding 5 µL 25% Tween-20 and incubated on ice for 10 min. Next, 25 µL samples were mixed with 30 µL High-Fidelity PCR Master Mix (NEB), 2.5 µL P5 primer (10 µM) and 2.5 µL P7 primer (10 µM) for PCR amplification. The PCR products were then purified before sequencing.

### RNA-seq data analysis

For bulk RNA-seq datasets, adaptor of all sequenced reads was first trimmed by Trim Galore with parameter “-illumina -paried”. Then the filtered reads were aligned to the mm10 reference genome by Hisat2 (version 2.0.0-b)^87^ with parameters “--no-mixed --no-discordant --dta-cufflinks --no-unal”. The aligned SAM files were then converted to BAM files with Samtools (version 1.14)^88^ for further analysis. Next, HTSeq-count (V0.12.4)^89^ was used to calculate the count of reads for each gene, with the main parameters set as: “-f bam -s no -r pos”. The reference genome was downloaded from the USCS Table Browser. To quantify gene expression, the read count for each gene was normalized using DESeq2 (V1.22.2).^90^ The differentially expressed genes from each group were filtered with following criteria: adjusted P-value < 0.05 and fold-change >2 or <0.5.

### ATAC-seq analysis

For ATAC-seq datasets, sequencing adaptors and low-quality reads were removed by Trim Galore with parameters: -q 20 --length 30 --paired. The clean reads were mapped to mouse reference genome (mm10) by bowtie2 (Version 2.5.1) with parameter: “-N 1 -L 25 -X 2000 -no-discordant”. After converting aligned SAM to BAM files using Samtools (Version 1.13), Picard (Version 2.26.4) was used for removing PCR duplications. Next, read counts were normalized by Reads Per Kilobase per Million mapped reads (RPKM) and converted to bigwig via bamCoverage (Version 3.5.1). The peaks were called by MACS2 (Version 2.2.7.1)^91^ with parameter “-t file -f BED -g mm -outdir OUTPUT -n NAME --nomodel --nolambda”. The peaks that overlap in more than 60% of the replicate samples in each group were kept for feature counts and subsequent analysis using DiffBind (V3.0.15).^92^ Peaks that differed significantly between groups were identified using DESeq2 with the following criteria: P-value < 0.05 and fold-change > 1.5 or < 0.67. Deeptools (Version 3.5.1)^93^ was used to visualize the read coverages at peak regions. Motif calling and peak annotation was performed using HOMER (V4.10) with all identified peaks as background. The genes within 5kb of analyzed peaks were regarded as peak-related genes.

### scRNA-seq data analysis

Paired reads from single-cell RNA seq were first aligned to mm10 reference genome with Cellranger --count (version 3.0.2) from 10X Genomics. Transcriptome annotation composed of protein coding genes was download and processed from Ensemble (released-102). To eliminate low-quality single cells, we filtered out single cells with feature numbers below 800 for freshly isolated HSCs (**Fig. 2**) and below 2,000 for short-term transplanted HSCs (**Fig. 4**). Additionally, cells with mitochondrial transcripts accounting for more than 5 percent were also removed. Then, the remaining single cells were processed and analyzed with Seurat (V4.0.2).^94^ The young and old single cell data is first merged and then normalized and scaled. Two thousand variable feature genes were identified and used for UMAP presentation. The pseudo time analysis was performed using Monocle (Version 2.22.0).^95^ The ordering genes are the marker genes of different cell types from pervious report.^50, 61^ The single cell GSEA analysis was performed using ClusterProfiler (Version 4.2.2).^96^

### Epigenetic age calculation

Paired sequencing files from RRBS were first merged and processed as a single file. Adaptors of all sequenced reads were trimmed by Trim Galore with parameters “-illumina”. Adaptor trimmed reads were mapped using Bismark (v0.20.0)^97^ using parameters “--fastq -L 30 -N 1 -- non_directional”. Methylated CpGs were interpreted by bismark_methylation_extractor with parameter “-s --no_overlap --report -bedGraph”. For base resolution 5mC level calculation, a cutoff of minimal 3× coverage was required for each CpG site. Epigenetic age was calculated based on the identified sites and parameters from a previous study.^98^

### Data accession

The dataset generated in this study is summarized in **Supplementary information, Table S7**. All data have been deposited to GEO with the accession number GSE233879.

## Acknowledgements

We thank Jiayi Li for her assistance in some experiments. We thank the support of Mouse Behavior Core of Harvard Medical School and its director Dr. Barbara Caldarone. This project was supported by the Milky Way Research Foundation and the Howard Hughes Medical Institute (HHMI). Y.Z. is an investigator of the HHMI. This article is subject to HHMI’s Open Access to Publications policy. HHMI lab heads have previously granted a nonexclusive CC BY 4.0 license to the public and a sublicensable license to HHMI in their research articles. Pursuant to those licenses, the author-accepted manuscript of this article can be made freely available under a CC BY 4.0 license immediately upon publication.

## Author Contributions

Y.Z. conceived and supervised the project; Y.W., W.Z. and Y.Z. designed the experiments; Y.W. performed the cell sorting, transplantation, FACS analysis, and physical function tests with help from H.T.V.; W.Z. and Y.W. performed the library preparation for bulk RNA-seq, scRNA-seq, RRBS and ATAC-seq; Y.W., W.Z. and C.Z. analyzed the single-cell RNA-seq data; Y.W. and W.Z. analyzed the bulk RNA-seq and RRBS data; Y.W. and C.Z. analyzed the ATAC-seq data; T.S. helped for complete blood counting analysis; Y.W., W.Z. and Y.Z. interpreted the data and wrote the manuscript with feedback from all authors.

---

**Fig. S1.**
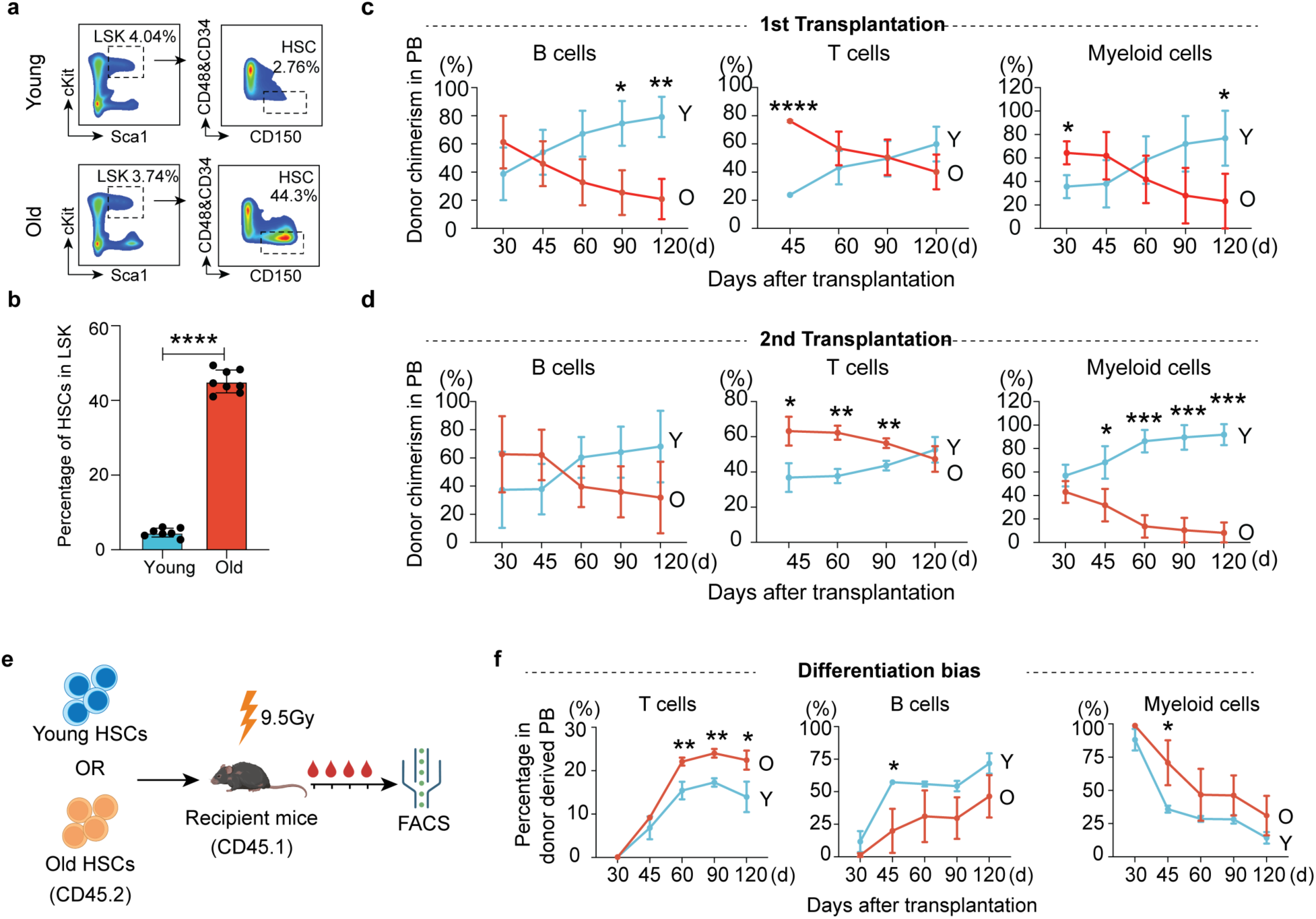
Aging-related functional decline of HSCs in mouse (related to Fig. 1). **a** The gating strategy for LT-HSCs sorting. The lineage^-^Sca-1^+^c-Kit^+^(LSK) was first gated, and LT-HSCs (CD48^-^CD34^-^CD150^+^) were gated in LSK population. **b** Bar graph showing the percentage of LT-HSCs in LSK HSPCs from young (2-3 months) and old (22-24 months) mice, *n* = 10 for young, *n* = 9 for old. **c-d** Peripheral blood chimerism of donor HSCs at different time after the first (**c**) and second (**d**) transplantation, *n* = 6 for the first and *n* = 3 for the second transplantation. **e** Diagram illustration of individual transplantation of young and old HSCs into young recipient mice for assessing differentiation bias. **f** Analysis of HSCs differentiation toward T, B and myeloid cells at different time after transplantation, *n* = 3. Mean ± SD, student t test, \**P*<0.05, ** *P*<0.01, *** *P*<0.001, **** *P*<0.0001. The graphic of the mouse in **e** was created with BioRender.

**Fig. S2.**
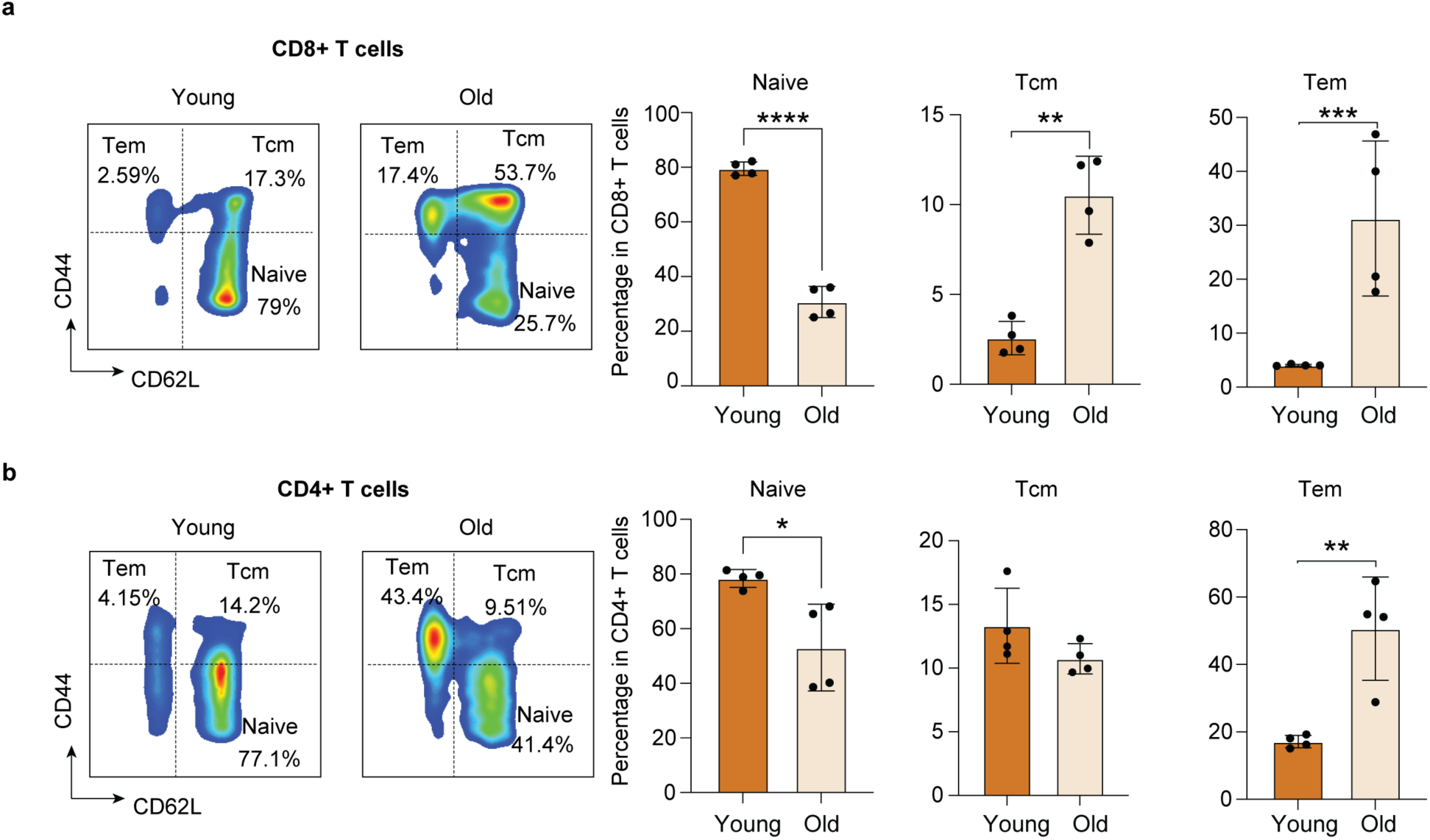
Immune profile alterations during aging in mice (related to Fig. 1). **a-b** Representative FACS analysis and bar plot showing the change in percentage of naïve, Tcm and Tem with aging in PB of mouse in CD8+ T cells (**a**) and CD4+ T cells (**b**). Mean ± SD, student t test, *n* = 4, * *P*<0.05, ** *P*<0.01, *** *P*<0.001, **** *P*<0.0001.

**Fig. S3.**
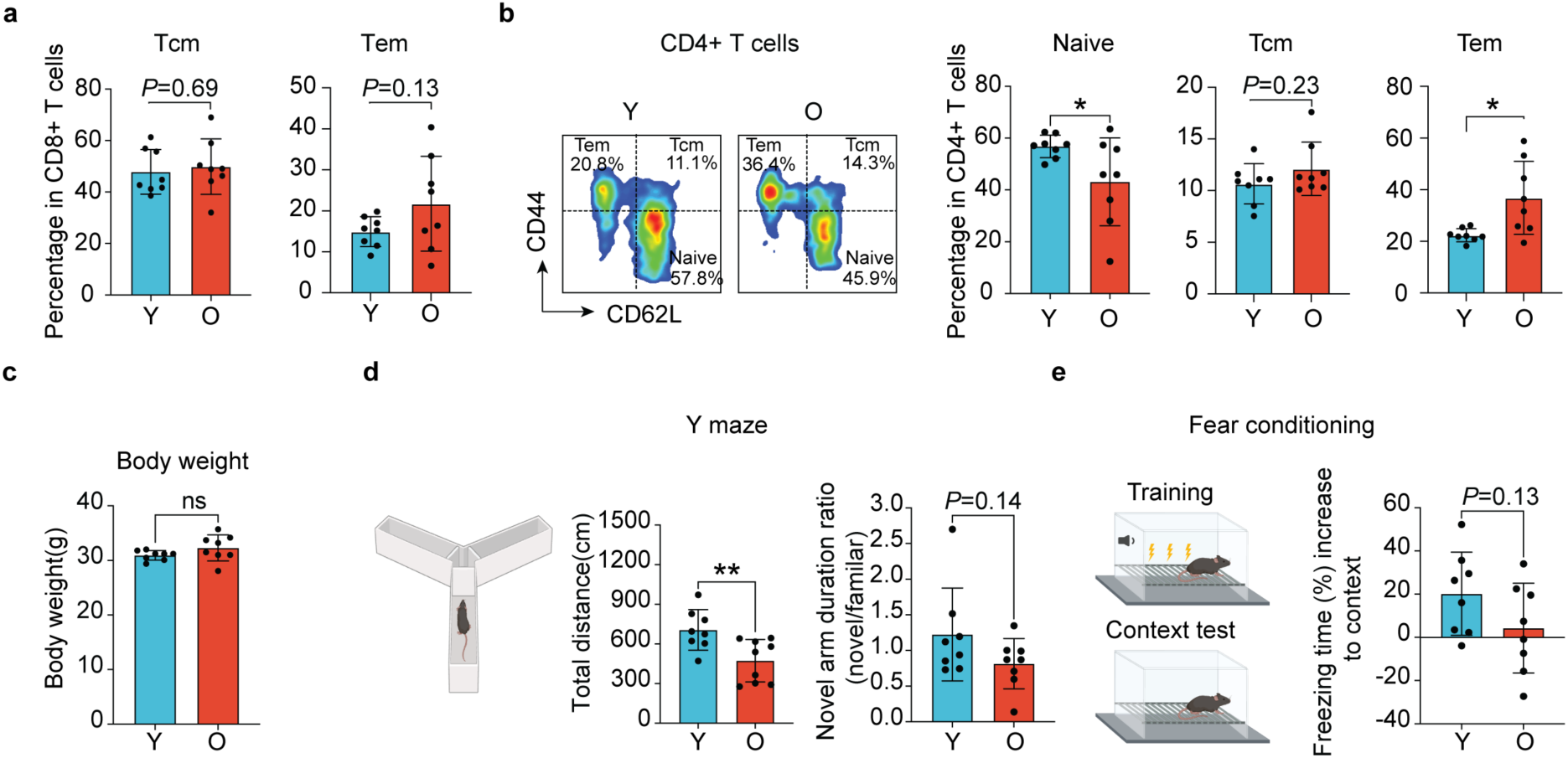
Transplantation of young HSCs alleviates aging phenotypes in old recipient mice (related to Fig. 1). **a** Bar plot showing the percentage of Tcm and Tem in CD8+ T cells from recipient mice, *n* = 8. **b** Representative FACS analysis and bar plot showing the percentage of naïve T cells, as well as Tcm and Tem in CD4+ T cells from recipient mice, *n* = 8. **c** Bar plot showing the body weight of recipient mice that received young or old HSCs, *n* = 8. **d** Bar plot showing the total moving distance and the ratio of duration in novel arm to familiar arm in Y maze test of recipient mice, *n* = 8. **e** Bar plot showing the results of context fear conditioning test. Compared to acclimation, the increased freezing time was calculated, *n* = 8. Mean ± SD, student t test, ** *P*<0.01, ns, not significant. The graphic of the mouse and equipment in **d** and **e** were created with BioRender.

**Fig. S4.**
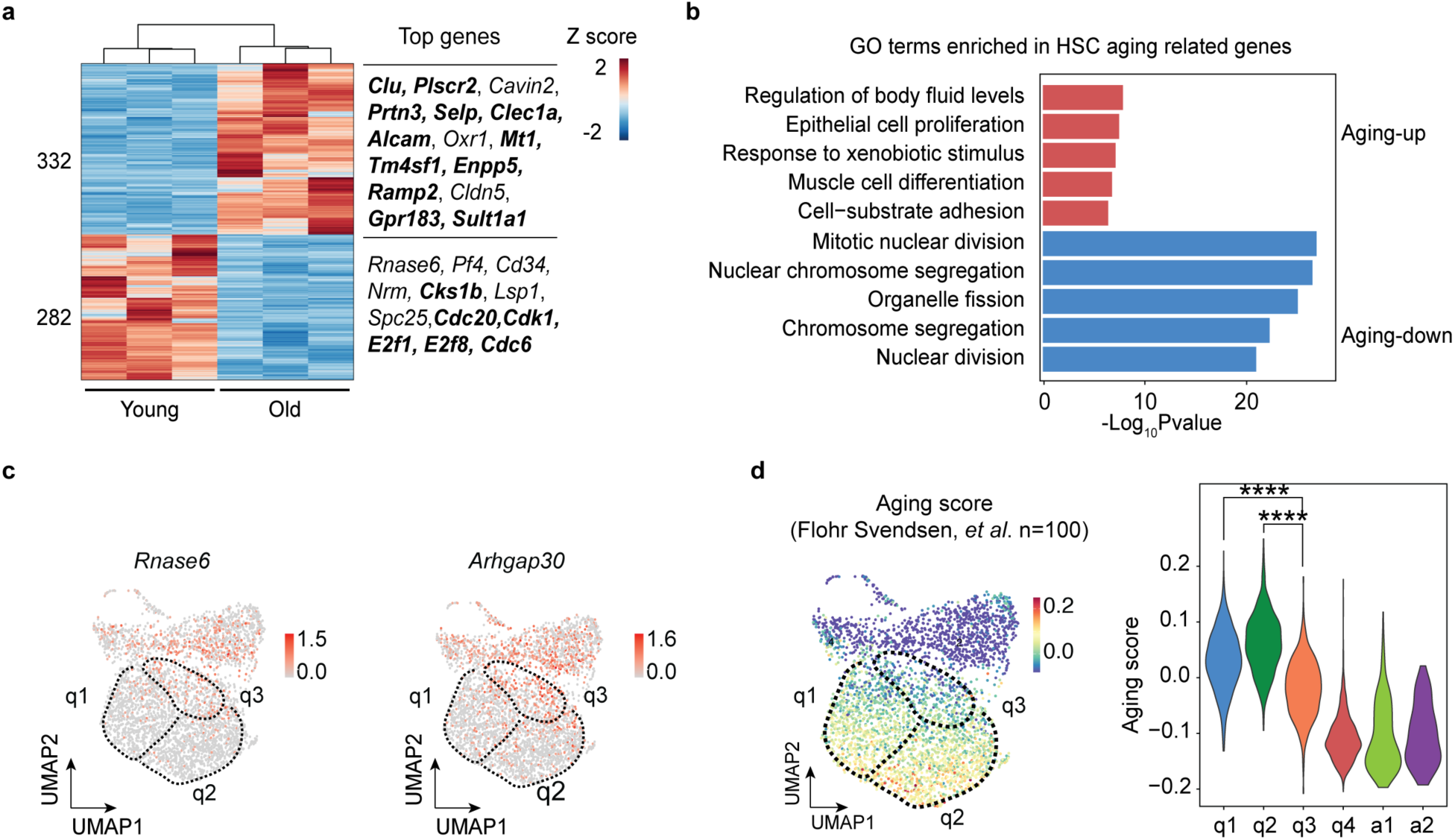
scRNA-seq reveals increased heterogeneity of old HSCs (related to Fig. 2). **a** Heatmap showing HSC aging related genes that were identified through bulk RNA-seq analysis. Well-known HSC marker genes in public datasets were highlighted. **b** Bar graph showing the enriched GO terms in HSC aging related up and down genes identified in (a). **c** UMAP showing the expression of the young HSC marker genes (*Rnase6* and *Arhgap30*) in single cells. **d** UMAP (left) showing the calculated aging score based on public HSC aging marker genes (Supplementary information, Table S3) in each single cell. Violin plot (right) showing the aging score of cells from different clusters. Two-sided unpaired Wilcoxon test, **** *P*<0.0001.

**Fig. S5.**
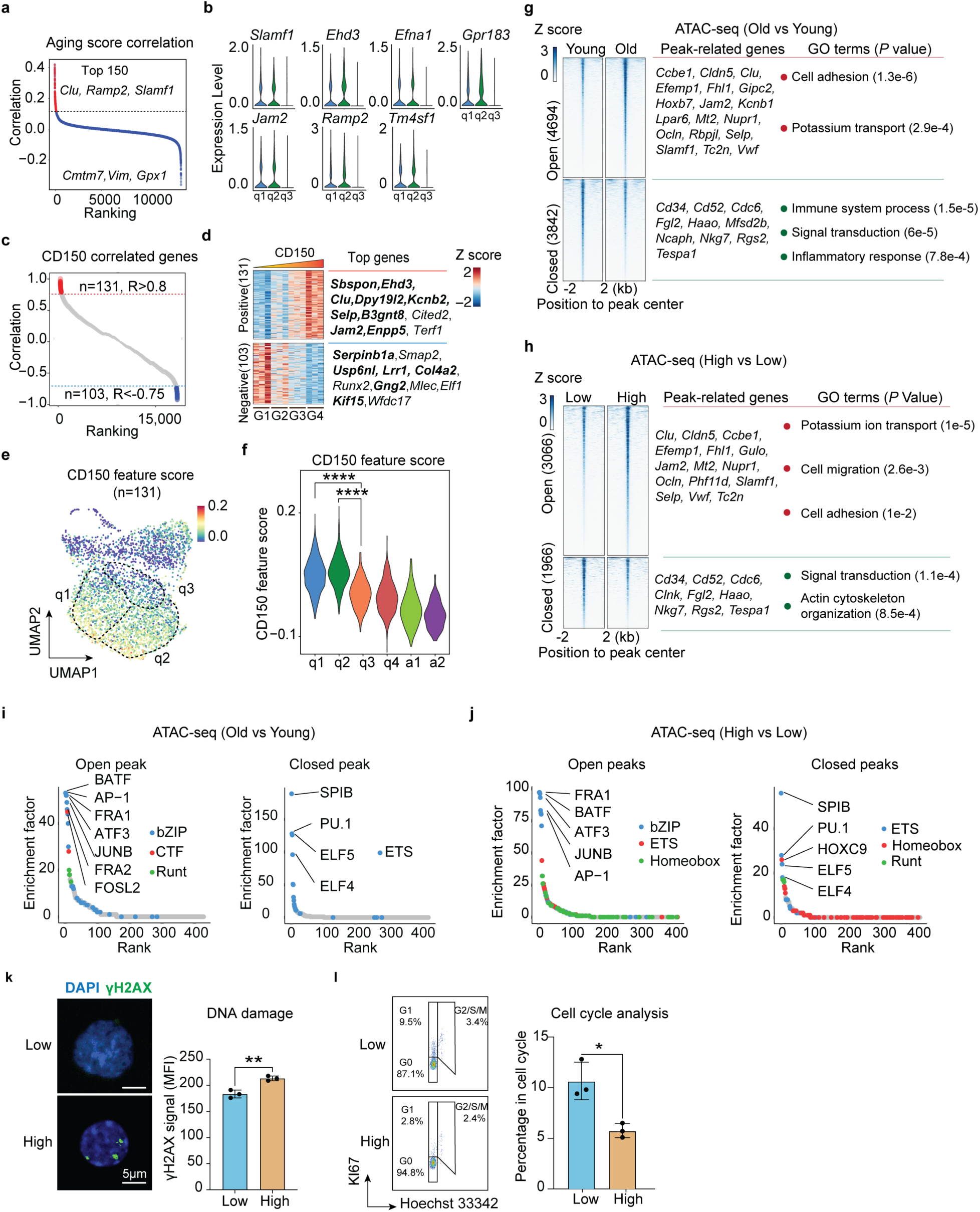
CD150 can serve as an aging heterogeneity marker of old HSCs (related to Fig. 3). **a** Dot plot showing ranking of genes based on the correlation of their expression and aging score in scRNA-seq. The top 150 highly ranked genes were highlighted. **b** Violin plot showing expression level of potential marker genes in cluster q1, q2 and q3 by scRNA-seq. **c** Dot plot showing the ranking of genes based on their correlation with CD150 levels in expression of the four cell groups in Figure 3C measured by bulk RNA-seq. A total of 131 positively correlated genes (R > 0.8) and 103 negatively correlated genes (R < −0.75) were identified as CD150 related genes. **d** Heatmap showing expression changes of CD150-related genes with ascending CD150 level based on bulk RNA-seq. For positive correlated genes, previously identified HSC aging marker genes are highlighted. For negative correlated genes, genes that are highly expressed in young HSCs are highlighted. **e-f** Feature plot (**e**) and violin plot (**f**) showing CD150 signature scores of the six cell clusters in scRNA-seq. **g** Heatmap showing aging-related open and closed peaks, as well as representative genes near these peaks (within 5 kb of the TSS) and the corresponding enriched GO terms followed by *P* value. **h** Heatmap showing the differential ATAC-seq peaks between old CD150^low^ HSCs and CD150^high^ HSC, as well as representative genes near these peaks (within 5 kb of the TSS) and the corresponding enriched GO terms, followed by P value. **i-j** Dot plot showing enriched TFs in aging-related open (left) and closed (right) peaks (**i**), and differential peaks between old CD150^low^ and CD150^high^ HSCs (**j**). The color of dot indicates different TF families. **k** Representative images and bar plot showing the level of γH2AX in old CD150^low^ and CD150^high^ HSCs, *n* = 3. **l** Left, representative FACS plot showing the percentage of old CD150^low^ and CD150^high^ HSCs in different cell cycle phases. Right, bar graph showing the percentage of old CD150^low^ and CD150^high^ HSCs in active cell cycle (G1 and G2/S/M), *n* = 3. For (g) and (h), Mean ± SD, student t test, \**P*<0.05, ** *P*<0.01. For (f), Two-sided unpaired Wilcoxon test, **** *P*<0.0001.

**Fig. S6.**
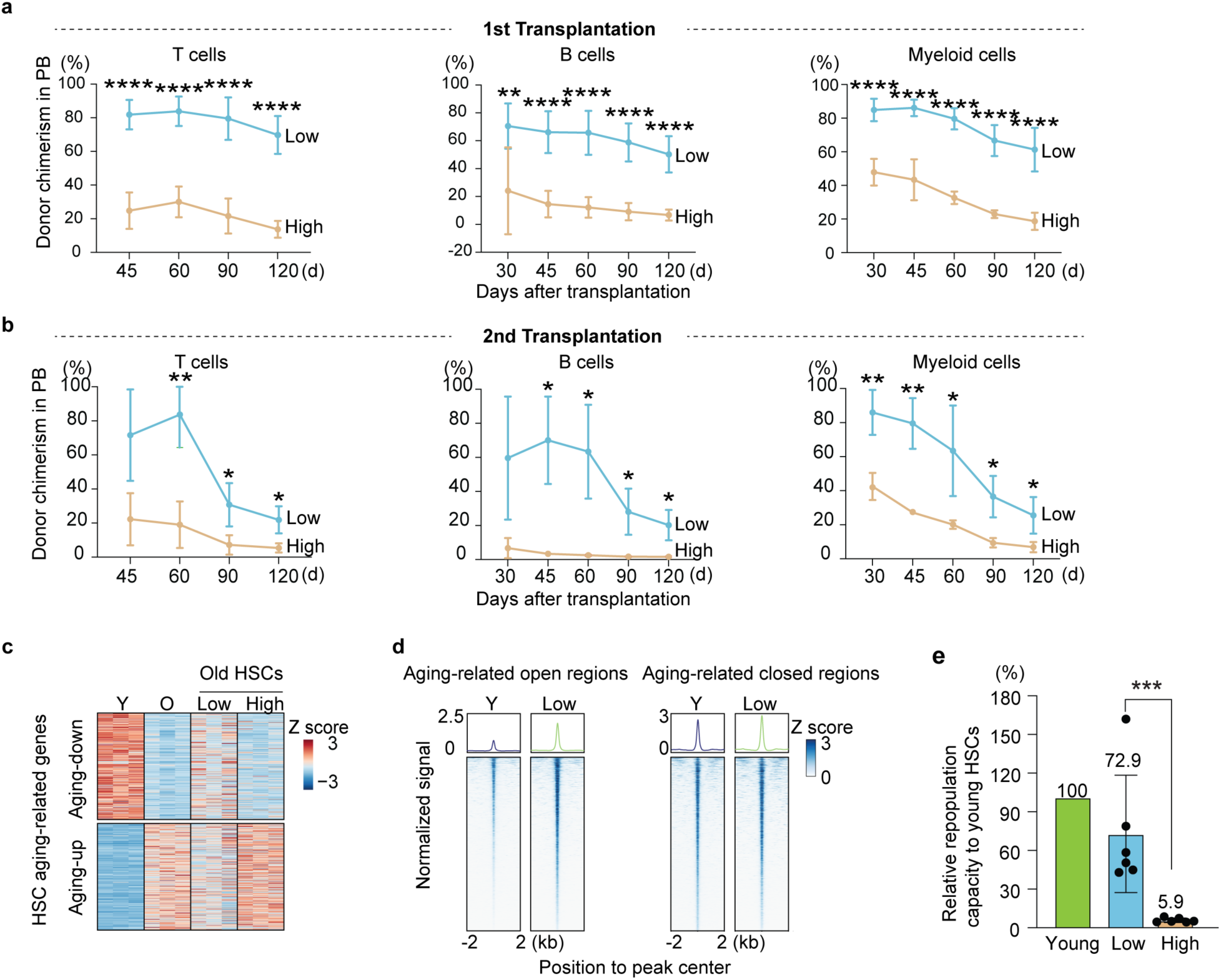
Comparison of old CD150^low^, CD150^high^, and young HSCs at the molecular and functional levels (related to Fig. 3). **a-b** The peripheral blood chimerism of donor HSCs at different time after transplantation in the first (**a**) and second (**b**) competitive transplantation. B, T, and myeloid cells were analyzed individually. Mean ± SD, student t test, *n* = 6 for the first and *n* = 3 for the second transplantation. **c** Heatmap showing the expression levels of aging-related up- and down-regulated genes in young and old HSCs, as well as old CD150^low^ and CD150^high^ HSCs. **d** Heatmap and bar plot illustrating ATAC-seq signals in aging-related open and closed chromatin peaks. **e** Bar plot showing the relative repopulation capacity of old CD150^low^ HSCs and CD150^high^ HSCs to young HSCs. For each sample, their relative functionality compared to young HSCs is calculated based on their contribution to PB, normalized by the number of transplanted cells. *n* = 6, Mean ± SD, student t test. \**P*<0.05, ** *P*<0.01, *** *P*<0.001, **** *P*<0.0001.

**Fig. S7.**
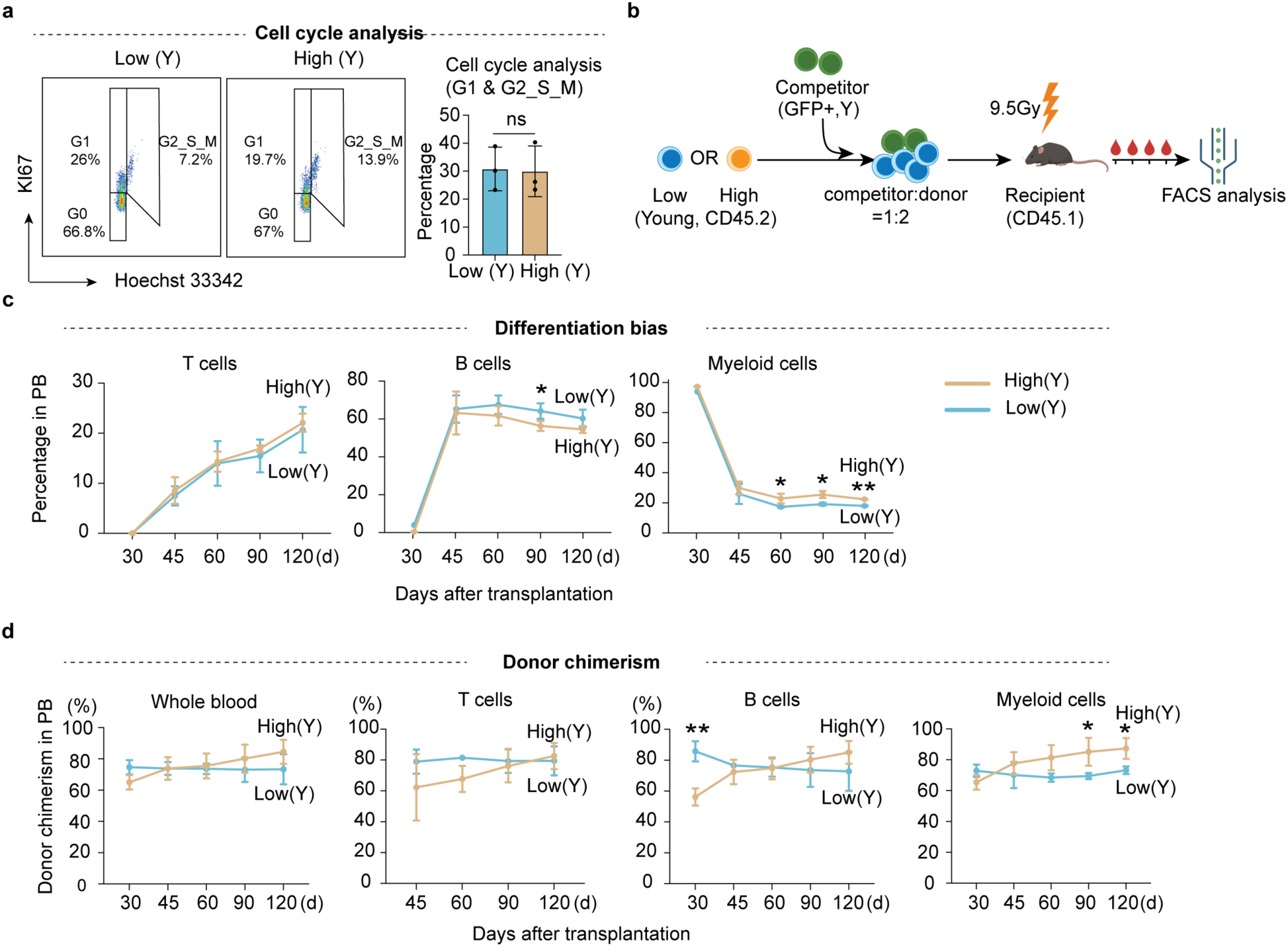
Comparable function of CD150^low^ and CD150^high^ HSCs from young mice (related to Fig. 3). **a** Left, representative FACS plot showing the percentage of young CD150^low^ (25% lowest) and CD150^high^ (25% highest) HSCs in different cell cycle phases. Right, bar graph showing the average percentage of young CD150^low^ and CD150^high^ HSCs in active cell cycle (G1 and G2/S/M), *n* = 3. **b** Diagram illustrating the competitive transplantation for evaluating the repopulating capacity of CD150^low^ and CD150^high^ HSCs from young mice. The ratio of competitor to donor HSCs to was 1:2 (300 competitor HSCs with 600 donor HSCs). **c-d** The differentiation bias (**c**) and peripheral blood chimerism (**d**) of donor HSCs at different time after transplantation. Whole blood, T, B, and myeloid cells were analyzed. *n* = 3, Mean ± SD, student t test, \**P*<0.05, \*\**P*<0.01, ns, not significant. The graphic of the mouse in **b** was created with BioRender.

**Fig. S8.**
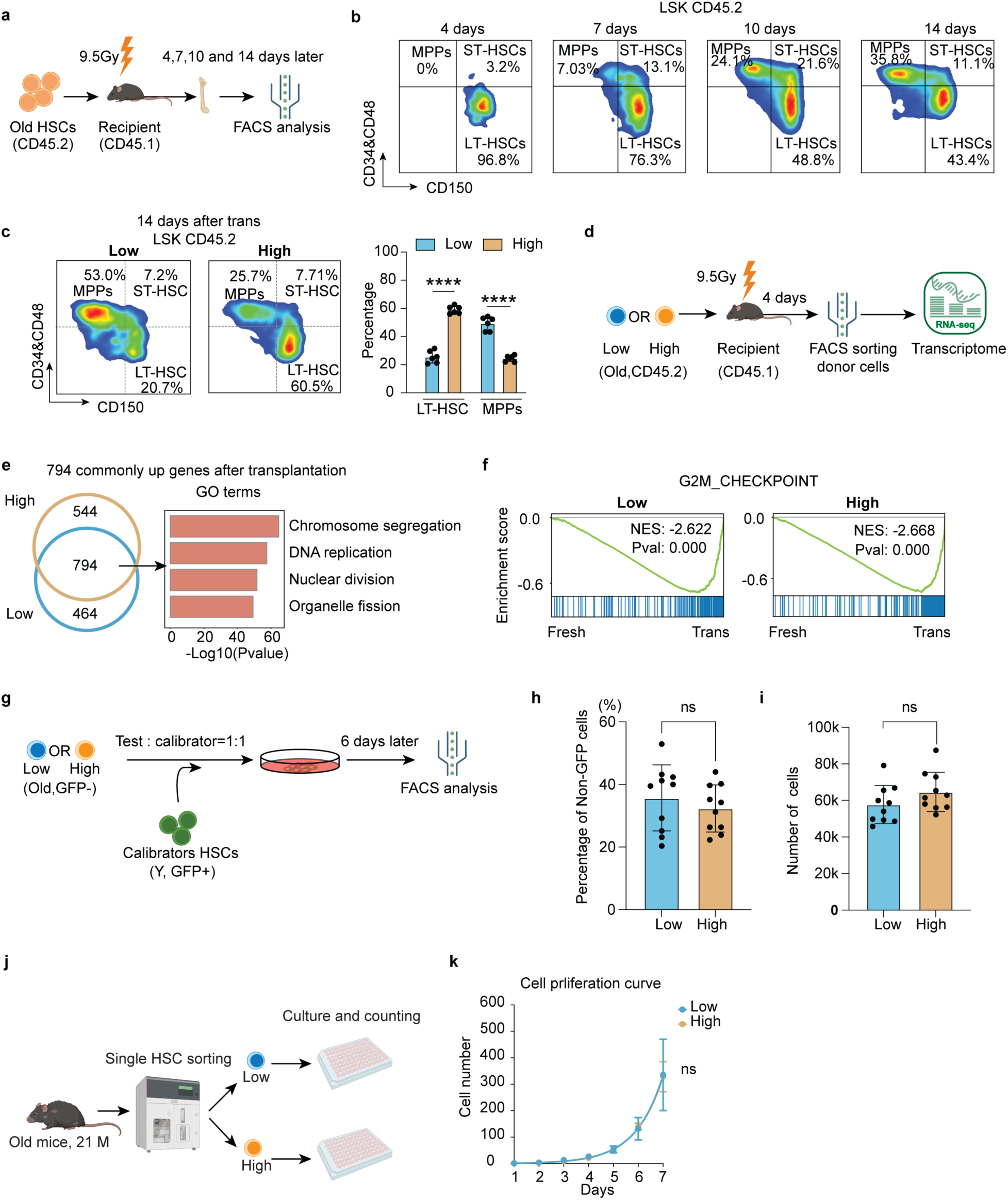
Differentiation, but not self-renewal, is a major defect of old CD150^high^ HSCs (related to Fig. 4). **a** Diagram of individual transplantation of old HSCs to study the differentiation trajectory after transplantation. The donor HSCs derived HSPCs in bone marrow were analyzed on days 4, 7, 10 and 14 after transplantation. **b** FACS analysis showing the differentiation trajectory of old HSCs after transplantation. On the 4^th^ day, most donor HSCs are still maintained as LT-HSCs. From 7^th^ days the LT-HSCs started to differentiate toward MPPs. **c** Representative FACS analysis of donor HSCs derived HSPCs (left) and quantification of different cell populations after transplantation (right) on day 14, *n* = 3. Mean ± SD, student t test. **d** Diagram of transplantation to evaluate the difference in activation between old CD150^low^ and CD150^high^ HSCs on day 4 after transplantation. The sorted donor HSCs were collected for transcriptome analysis, *n* = 3. **e** Venn diagram showing the genes that were commonly activated in old CD150^low^ and CD150^high^ HSCs compared to freshly isolated old CD150^low^ and CD150^high^ HSCs 4 days after transplantation (left). Bar graph showing enriched GO terms of the 794 commonly activated genes (right). **f** GSEA analysis showing that transplanted HSCs highly express cell cycle related genes in both CD150^low^ (left) and CD150^high^ (right) HSCs when compared with freshly isolated HSCs. **g** Diagram showing the experimental design for examining HSCs proliferation *in vitro*. In each well, 50 GFP+ cell and 50 old CD150^low^ or CD150^high^ HSCs were co-cultured. The percentage and absolute number of old CD150^low^ and CD150^high^ HSCs were quantified 6 days after culture. **h** Bar graph showing comparable proliferation rate of old CD150^low^ and CD150^high^ HSCs 6 days after culture. The percentage of non-GFP HSCs was shown, *n* = 10. Mean ± SD, student t test. **i** Bar graph showing the absolute number of old CD150^low^ and CD150^high^ HSCs 6 days after culture, *n* = 10. Mean ± SD, student t test. **j** Diagram showing the workflow for evaluating the proliferation capacity of old CD150^low^ and CD150^high^ HSCs. **k** Cell proliferation curve showing the cell number change with time (1-7 days), *n* = 8, student t test, Mean ± SEM. \*\**P*<0.01, *** *P*<0.001, ns, not significant. The graphic of the mouse and equipment in **a**, **d** and **j** were created with BioRender.

**Fig. S9.**
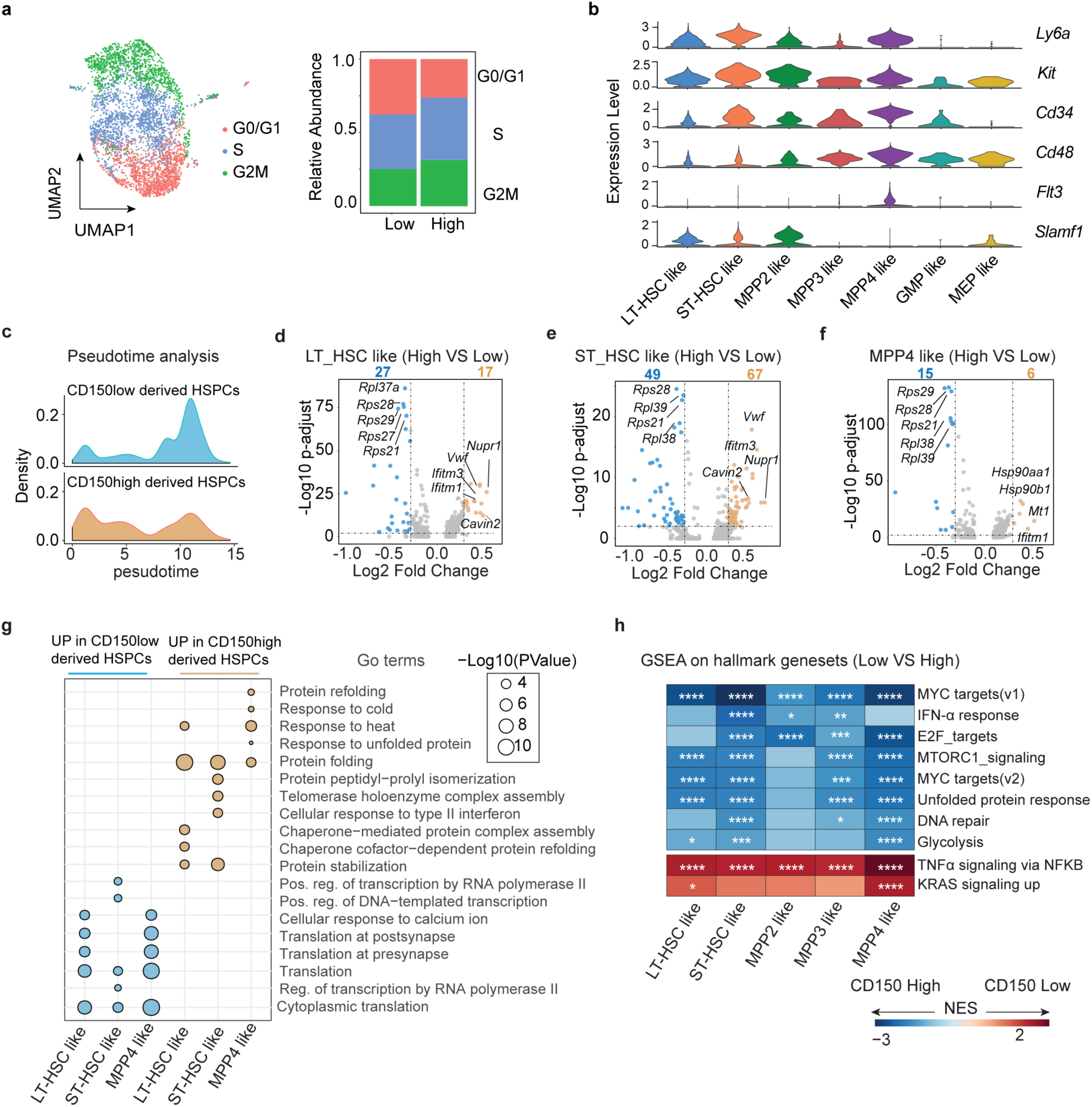
Old CD150^high^ HSCs are defective in the LT-HSCs to ST-HSCs transition (related to Fig. 4). **a** UMAP and bar graph showing the distribution and percentages of cells in different cell cycle phases of HSPCs from CD150^low^ and CD150^high^ HSCs 14 days after transplantation. **b** Violin plot showing the expression pattern of cell type marker genes, including *Ly6a*, *Kit*, *Cd34*, *Cd48*, *Flt3* and *Slamf1* (CD150). **c** Density plot showing the relative abundance of cells at different pseudo time stage. **d-f** Volcano plot showing the differentially expressed genes between old CD150^low^ and CD150^high^ HSCs derived HSPCs, including LT-HSCs-like (**d**), ST-HSCs-like (**e**) and MPP4-like (**f**) cells. **g** Dot plot showing the enriched GO terms in differentially expressed genes between old CD150^low^ and CD150^high^ HSCs derived HSPCs. The size of dot indicates the significance of enrichment. **h** Heatmap displaying the GSEA comparison between transplanted old CD150^low^ and CD150^high^ HSC-derived HPSC cell types, with red indicating activated pathways in CD150^low^ HSCs derived cell types and blue indicating activated pathways in old CD150^high^ HSCs derived cell types. \**P*<0.05, ** *P*<0.01, *** *P*<0.001, **** *P*<0.0001.

**Fig. S10.**
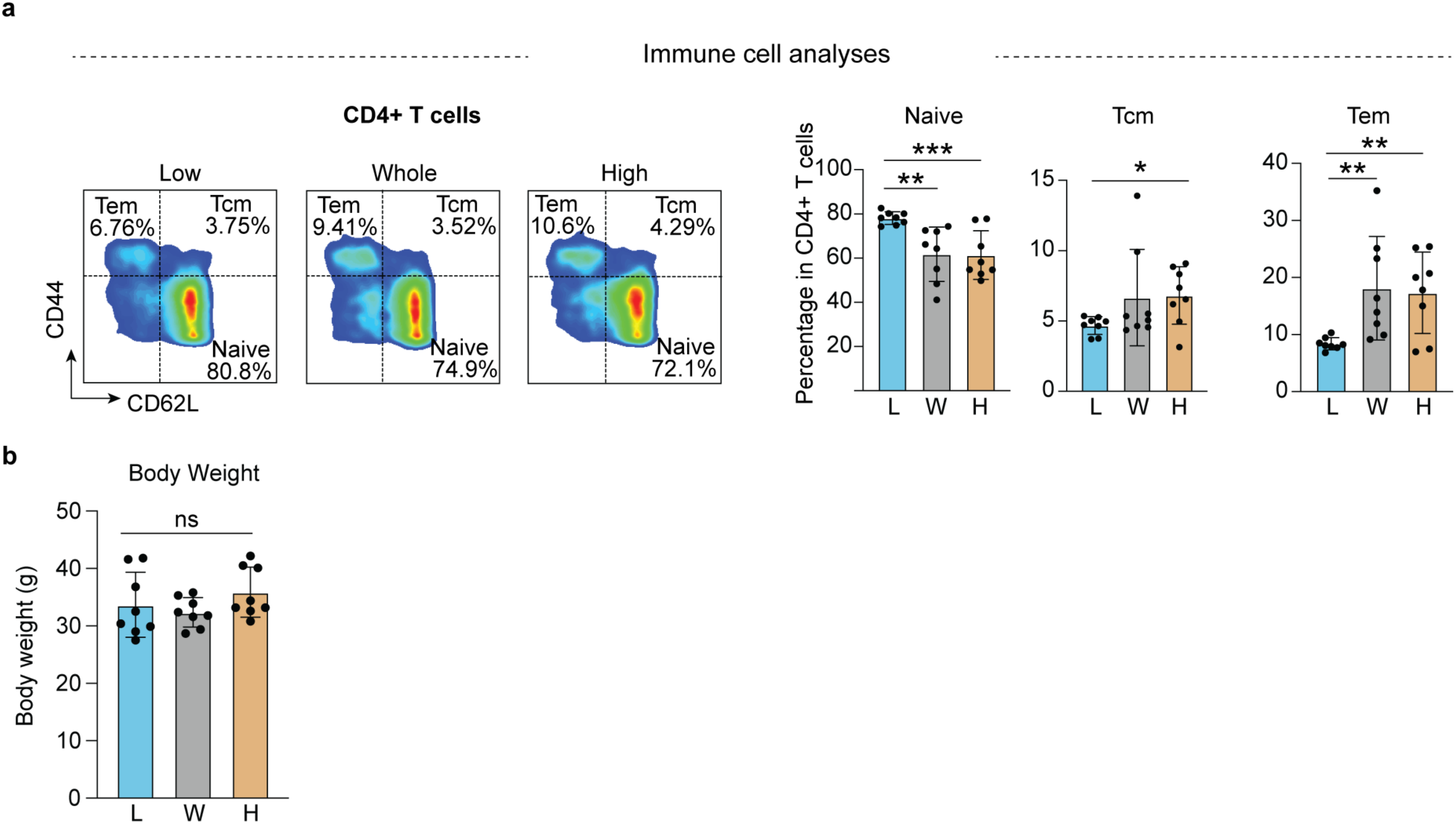
Transplantation of “younger” subset old HSCs attenuates aging phenotypes of old mice (related to Fig. 5). **a** Representative FACS (left) analysis of naïve T cells, Tcm and Tem ratio in CD4 positive T cells of mice from different groups and their quantification (right, bar graphs). **b** Bar graph showing body weight of recipient mice from different groups. Mean ± SD, one-way ANOVA, *n* = 8, \**P*<0.05, ** *P*<0.01, *** *P*<0.001, ns, not significant.

**Fig. S11.**
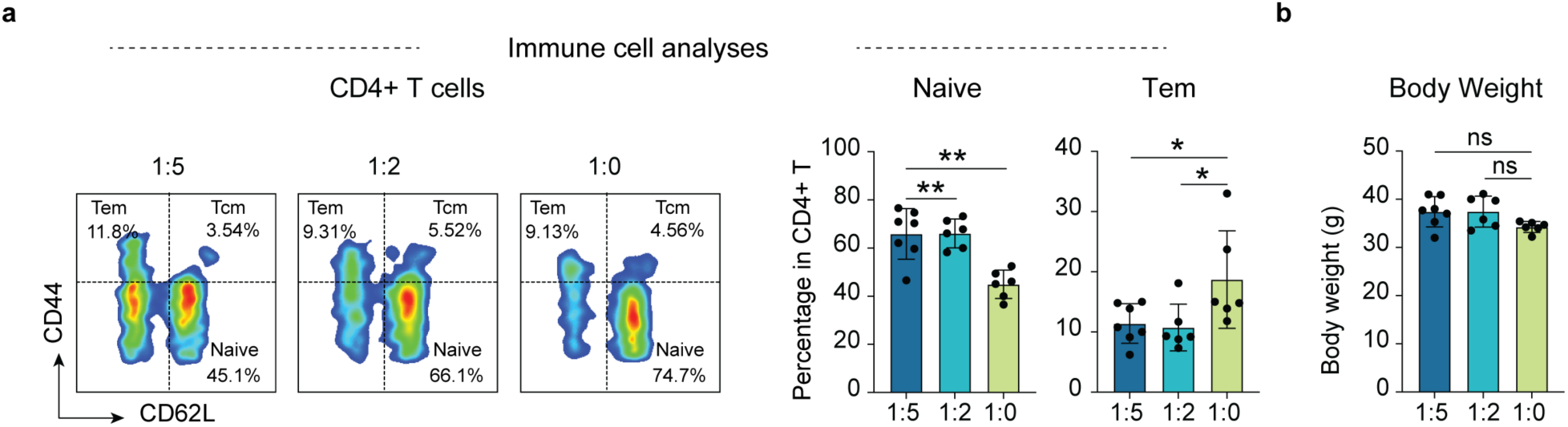
Reducing dysfunctional HSCs ameliorates aging phenotypes in old mice post transplantation (related to Fig. 6). **a** Representative FACS (left) analysis of naïve T cells and Tem ratio in CD4 positive T cells of mice from the different groups and their quantification (right, bar graphs). **b** Bar graph showing body weight of recipient mice from different groups. n=6 for 1:5 and 1:2 group, *n* = 7 for 1:0 group; Mean ± SD, one-way ANOVA, \**P*<0.05, ** *P*<0.01, ns, not significant.

